# BCAA catabolism drives adipogenesis via an intermediate metabolite and promotes subcutaneous adipose tissue expansion during obesity

**DOI:** 10.1101/2022.08.18.504380

**Authors:** Jing Shao, Yunxia Liu, Xuejiao Zhang, Le Shu, Jiayu Yu, Sa Yang, Chen Gao, Chenma Wang, Nancy Cao, Meiyi Zhou, Rui Chi, Mengping Chen, Chunliang Liu, Ji Wang, Weiping Zhang, Ruixin Liu, Jiqiu Wang, Weiqing Wang, Guang Ning, Xia Yang, Yibin Wang, Haipeng Sun

## Abstract

Branched-chain amino acids (BCAAs, including leucine, isoleucine, and valine) have emerged as major players in metabolic health and diseases, but the underlying mechanisms remain obscure. Here, we report that BCAA catabolism drives adipogenesis via an intermediate metabolite of leucine and promotes subcutaneous white adipose tissue (sWAT) expansion during obesity. Genetic analyses of humans and mice reveal that the BCAA catabolic pathway in WAT is strongly correlated with adipose physiology and obesity traits. Altering BCAA catabolism in mature adipocytes exerts minor effects on adiposity in mice. However, enhancing adipose BCAA catabolism via FABP4-Cre-mediated *Bckdk* deletion promotes diet-induced obesity while blocking adipose BCAA catabolism through *Bckdha* ablation does the opposite. Intriguingly, the catabolism of BCAAs elicits fat depot-specific responses and promotes sWAT extension and adipogenesis in a cell-autonomous manner. Mechanistically, BCAA catabolism drives adipocyte differentiation via an intermediate metabolite of leucine, which activates mTORC1 and polyamine synthesis from methionine to promote the expression of adipogenic master regulators. Together, these results demonstrate that BCAA catabolism promotes adipogenesis and sWAT expansion during obesity. The crosstalk between leucine and methionine metabolism driven by the catabolic intermediate highlights an unexpected regulatory role of amino acids in metabolic health and diseases.

## INTRODUCTION

In addition to lipid and glucose, amino acids, particularly the branched-chain amino acids (BCAAs, including leucine, isoleucine, and valine), have emerged as major players in metabolic health and diseases. Disrupted BCAA homeostasis has been strongly correlated with obesity and obesity-associated disorders with potential causal roles ^1–9^. However, it is noteworthy that the detrimental effects of BCAA metabolic dysregulation appear to be tightly associated with the pathological obesity. In lean mice, blocking BCAA catabolism leads to attenuated obesity and enhanced insulin sensitivity ^10, 11^. These paradoxical impacts of BCAA metabolic dysregulation are in line with the different or even opposite metabolic effects of dietary BCAA intake ^1, 3, 12–14^. The mechanisms underlying these metabolic impacts remain to be fully determined.

BCAA homeostasis is tightly controlled by their catabolic pathway that consists of dozens of enzymes and catabolic intermediates prior to the end-products ^1, 4^. The first two steps of BCAA catabolism are common to all three BCAAs. BCAA transaminase (BCAT) catalyzes the initial transamination to produce branched chain keto acids (BCKAs). The second and rate-limiting step is catalyzed by BCKA dehydrogenase (BCKD) complex. BCKD kinase (BCKDK) phosphorylates the BCKD E1α subunit (BCKDE1α) to inhibit BCKD activity. After the BCKD step, the individual BCAA catabolic pathways diverge with different enzymes and processes. BCAA intake and catabolism coordinate to determine the abundances of BCAAs and their catabolites, many of which exert important metabolic functions ^4^. BCAA *per se* modulates food intake, insulin sensitivity, energy expenditure, and functions of hormones including insulin, leptin, and glucagon, largely via their signaling function ^1^. A valine catabolic intermediate, 3-Hydroxyisobutyrate (HIB), regulates vascular fatty acid uptake and insulin sensitivity in skeletal muscle ^15^. The leucine-derived acetyl-CoA can be an activator of mTORC1 signaling or a major source of priming substrates for *de novo* synthesis of fatty acids ^16–18^. Thus, alterations of BCAA catabolism may affect lipid and glucose metabolism via both BCAAs and their metabolites.

The catabolism of BCAA occurs in multiple organs ^1, 19, 20^. Both white adipose tissue (WAT) and brown adipose tissue (BAT) are critical regulators of systemic BCAA homeostasis. BAT dissipates energy as heat to maintaining body temperature and improves glucose and lipid homeostasis. A recent study shows that BCAA catabolism in BAT contributes to fuel thermogenesis and resist diet-induced obesity and insulin resistance in mice ^19^. WAT stores excess energy as lipid and plays a beneficial role in maintaining metabolic homeostasis during the onset of obesity. However, in the context of pathological obesity, WAT demonstrates a detrimental role to promote complications such as insulin resistance. BCAA catabolism in WAT is dramatically suppressed in obese animals and humans, which contributes to the elevation of plasma BCAA levels ^1, 20^. Intriguingly, a recent study shows that mice with adipose tissue depletion of BCAA catabolism are resistant to diet-induced obesity due to increased subcutaneous WAT (sWAT) browning and thermogenesis ^21^. Furthermore, BCAA catabolism is essential for 3T3-L1 adipocytes differentiation by providing substrates for lipid synthesis *in vitro* ^17, 22–24^. On the other hand, a recent study elegantly demonstrates that WAT is not a major tissue for BCAA disposal in mice ^20^. The function of BCAA catabolism in WAT and its impact on obesity and associated disorders remain to be fully elucidated.

Here, we demonstrate that the BCAA catabolic pathway in WAT is genetically linked with WAT physiology and obesity traits in humans and mice. Furthermore, adipose BCAA catabolism promotes diet-induced obesity and sWAT expansion in a depot-specific manner. We also found that BCAA catabolism promotes obesogenic adipogenesis cell-autonomously in sWAT. A leucine catabolic intermediate activates mTORC1 and drives a metabolic network involving methionine and polyamine to promote the expression of adipogenic master regulators. These results reveal a previously unrecognized regulatory role of BCAA catabolism in adipogenesis, WAT physiology, and the development of obesity.

## RESULTS

### Adipose BCAA catabolic pathway is genetically associated with obesity traits in humans and mice

In our previous evaluation of the association between biochemical pathways and metabolic traits in the Hybrid Mouse Diversity Panel (HMDP) consisting of a population of over 100 inbred mouse strains, the BCAA catabolic pathway in WAT exhibited strong correlation with HFD-induced insulin resistance ^5^. Further analysis of the HMDP data revealed a significant correlation with body weight, adiposity, and fat mass for the majority of the BCAA catabolic genes expressed in WAT in mice (**Fig. 1A**). The BCAA catabolic genes as a group also demonstrated stronger correlation strength compared to the background genes (**Fig. 1B**). Importantly, the correlation strength between BCAA catabolic genes and the metabolic traits escalated upon high-fat diet (HFD) challenge (**Fig. 1A**), while other non-BCAA amino acid metabolic genes did not show similar sensitivity (Fig. S1). The genetic links observed in the HMDP mouse population strongly suggest the physiological importance of BCAA catabolism in adipose tissue and the development of obesity.

**Fig. 1.**
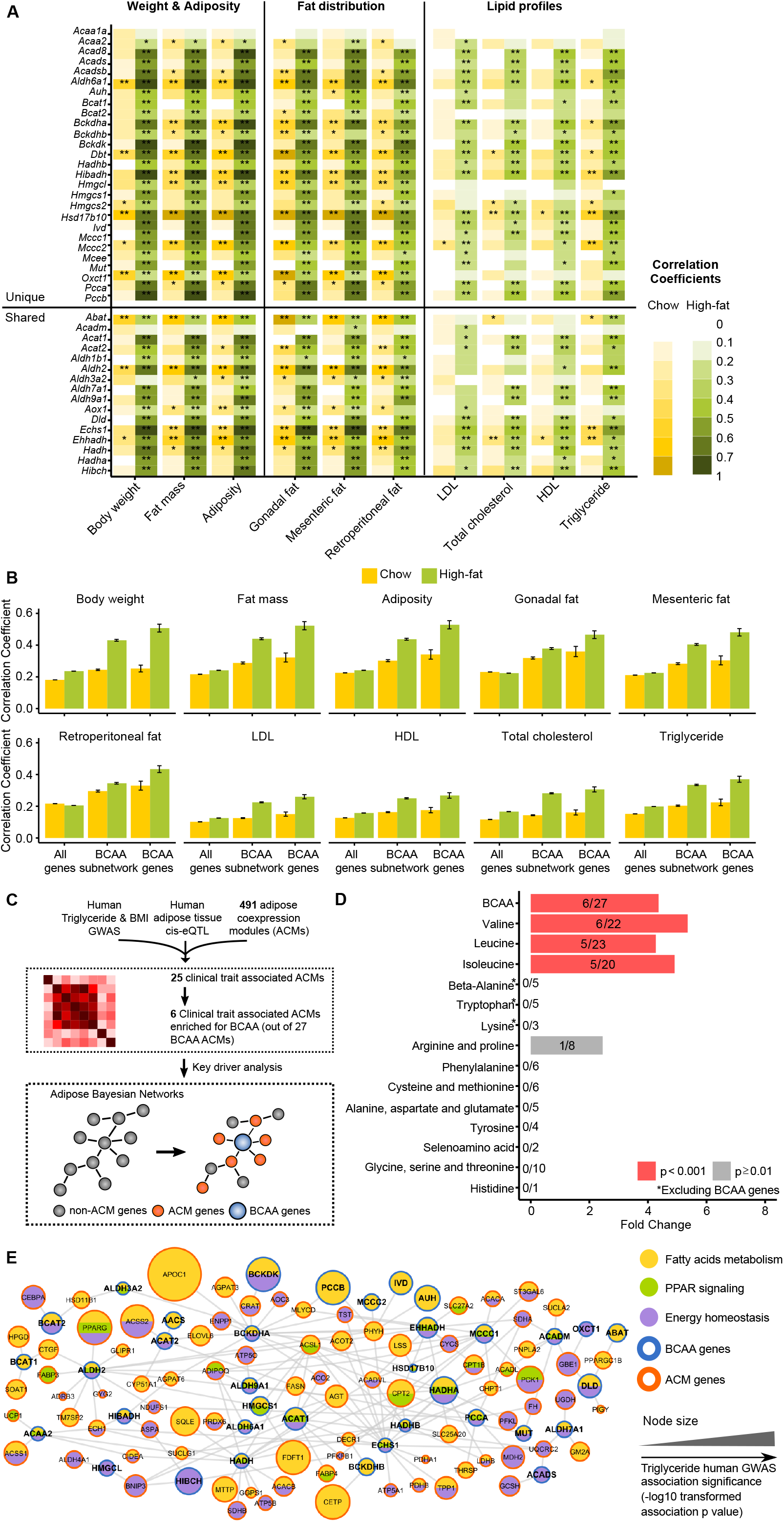
Genetic association between adipose BCAA catabolic pathway and obesity traits in the Hybrid Mouse Diversity Panel (HMDP) and human GWAS studies. (A) Gene-trait correlation for BCAA catabolic genes in the adipose tissue from HMDP mice fed with chow diet and high-fat diet. *passing Bonferroni p < 0.05 corrected for the number of BCAA catabolic genes; **passing Bonferroni p < 0.05 corrected for number of BCAA catabolic genes and the number of traits. (B) Comparison of average correlation coefficients with traits for background genes, genes in BCAA subnetwork and BCAA catabolic genes in adipose tissue from HMDP mice fed with chow diet or high-fat diet. Error bars indicate standard errors. (C) Design of integrative genomics analysis of human GWAS studies of obesity traits. Human GWAS results for plasma triglyceride level and BMI were first integrated with human adipose cis-eQTL and adipose coexpression modules (ACMs) to identify obesity-associated ACMs, which were subsequently annotated with biological pathways including BCAA catabolism. Next, BCAA ACMs were mapped to adipose Bayesian networks to test whether BCAA genes play regulatory role on these obesity-associated ACMs. (D) Association of ACMs involving various amino acids metabolic pathways with plasma triglyceride level and BMI. The numbers in each bar represent (the number of obesity-associated ACMs with enrichment of the particular amino acid pathway)/(the total number of ACMs annotated with the same amino acid pathway). Fisher’s exact test was used to assess whether the obesity-associated ACMs (total 25) were enriched for those annotated with any particular amino acid pathway. (E) Relationship between BCAA genes and the co-regulated metabolic pathways in an adipose Bayesian network. Edge colors indicate the metabolic functions of genes connected to BCAA genes. The size of each node is proportional to triglyceride association strength (-log10 transformed p value) in human GWAS.

We also employed unbiased genomics analysis to investigate the potential causal links between biological pathways and obese traits in human genome-wide association studies (GWAS) (**Fig. 1C**). A total of 25 out 491 adipose co-expression modules (ACMs) were found to be significantly enriched for genetic variants associated with plasma triglyceride (TG) and/or body mass index (BMI) (Supplementary Table 1). These 25 modules implicated immune pathways, TCA cycle, oxidative phosphorylation, etc. Importantly, 6 of these 25 modules were enriched for genes in the BCAA catabolic pathway, representing a 4.4-fold over-representation of the BCAA catabolic pathway in the TG/BMI ACMs while non-BCAA pathways lack such enrichment (**Fig. 1D**). After applying a network analysis to the TG/BMI ACMs enriched for BCAA metabolic genes using an adipose Bayesian network, we found BCAA catabolic genes coordinated the activities of genes that function in fatty acid metabolism, PPAR signaling, and energy homeostasis (**Fig. 1E**). These results suggest a role of BCAA catabolic pathway in affecting adipose tissue physiology and obesity traits in human.

### Enhancing adipose BCAA catabolism promotes diet-induced obesity in *Ap2*-*Bckdk*-AKO mice

Next, we generated conditional knockout mouse lines to analyze the impacts of enhancing BCAA catabolism on adipose physiology. Several Cre transgenic mouse models have been used to target adipose tissues, each with their own limitations ^25–29^. We generated mice carrying the floxed *Bckdk* allele containing two loxP sites flanking exons 2-8 (*Bckdk ^f/f^*) and cross them with either *Fabp4*-Cre (*Ap2*-*Bckdk*-AKO) (**Fig. S2A and S2B**) or *Adipoq*-Cre (*Adq*-*Bckdk*-AKO) (**Fig. S2O**) transgenic mice, respectively ^30^. These mouse lines provide tools to analyze the metabolic impacts of the enhanced BCAA catabolism in either mature adipocytes in *Adq*-*Bckdk*-AKO mice or numerous types of adipose cells in *Ap2*-*Bckdk*-AKO mice, in spite of that the *Fabp4*-Cre mediated gene deletion could occur in non-adipose tissues ^27^.

In *Ap2*-*Bckdk*-AKO mice, BCKDK protein expression and BCKDE1α phosphorylation were significantly reduced in the epididymal WAT (eWAT, representative of visceral WAT, vWAT), inguinal WAT (iWAT, representative of sWAT), and BAT, but not in other major tissues (**Fig. S2C**). Metabolomic analysis demonstrated elevated abundances of BCKD products in the sWAT of *Ap2*-*Bckdk*-AKO mice (**Fig. S2D**), further supporting the enhanced BCAA catabolism.

On normal chow diet (NCD), the body weight, lean mass, fat mass, organ weight, and glucose and insulin tolerance of *Ap2*-*Bckdk*-AKO mice were not different from those of control littermates (**Fig. S2E-S2I**). However, in response to HFD feeding, the body weight gain of *Ap2*-*Bckdk*-AKO mice was significantly higher than that of littermate mice (**Fig. 2A**). After 6-week HFD feeding, a significant increase in fat but not lean content was observed in *Ap2*-*Bckdk*-AKO mice (**Fig. 2B**), accompanied with greater mass of eWAT, iWAT, and BAT (**Fig. 2C**), enlarged adipocytes (**Fig. 2D**), and impaired glucose clearance and insulin sensitivity (**Fig. 2E** and **2F**). The oxygen consumption and heat production were lower in HFD-fed *Ap2*-*Bckdk*-AKO mice compared with their littermates (**Fig. S2J-S2L**), likely contributing to the accelerated obesity. Physical activity of HFD-fed *Ap2*-*Bckdk*-AKO mice were comparable to those of *Bckdk^f/f^* mice (**Fig. S2M**). The food intake did not show difference (**Fig. S2N**).

**Fig. 2.**
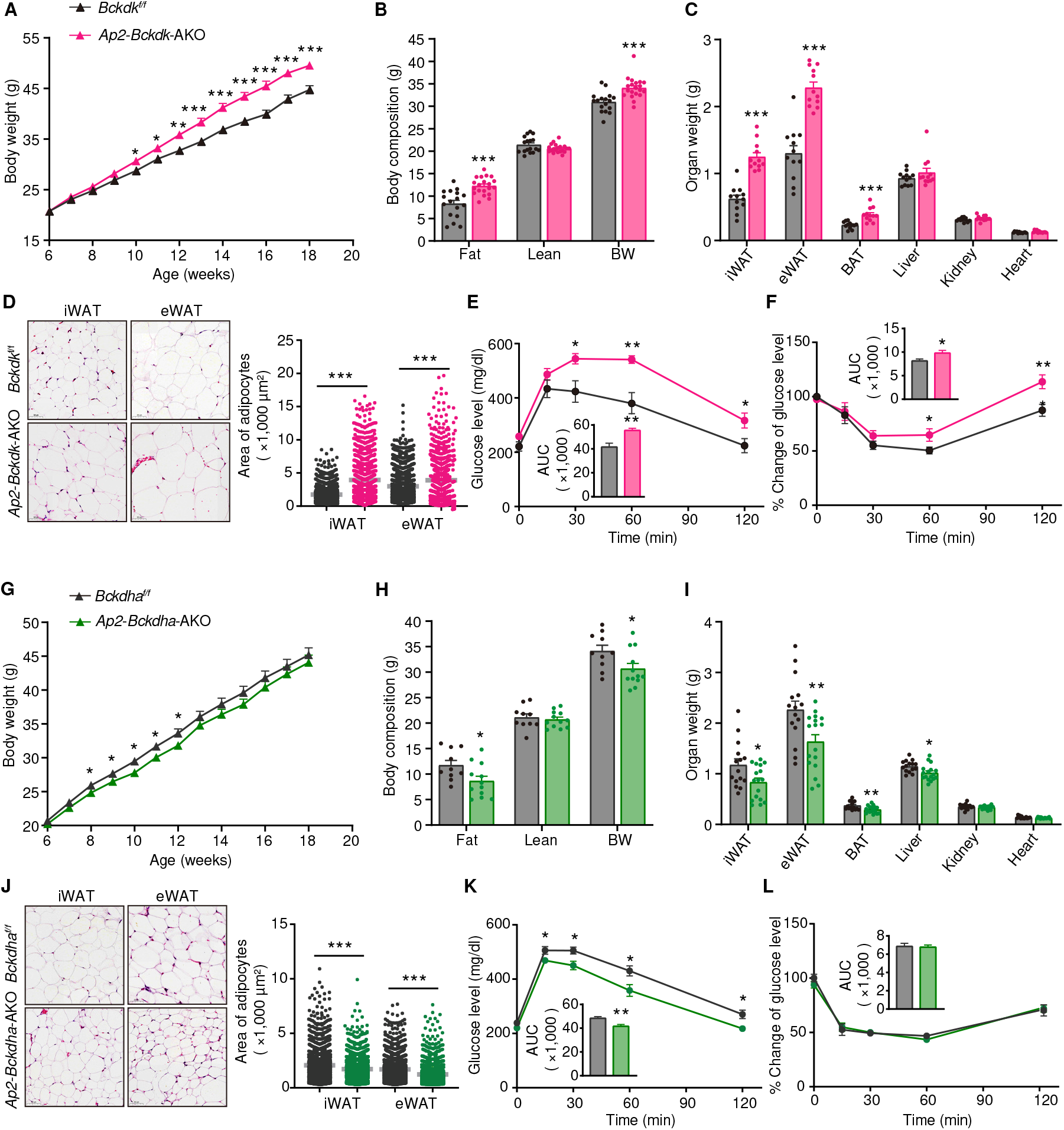
Manipulating adipose BCAA catabolism affects the development of diet-induced obesity in mice. (A-F) *Bckdk ^f/f^* and *Ap2*-*Bckdk*-AKO mice were fed with 60% HFD for 12 weeks (A) and for 6 weeks (B-F). A, Body weight (n=17-26 per group). B, Body composition (n=17-20 per group). C, Organ weight (n=12 per group). D, Images of iWAT and eWAT. Scale bars, 50 μm. Quantification of adipocytes surface areas (right, n=1025-1567 cells per group). E, Glucose tolerance test (GTT) (n=6-7 per group) and the area under the curve (AUC). F, Insulin tolerance test (ITT) (n=8-9 per group) and the AUC. (G-L) *Bckdha ^f/f^* and *Ap2*-*Bckdha*-AKO mice were fed with 60% HFD for 12 weeks (G) and for 6 weeks (H-J). G, Body weight (n=20-21 per group). H, Body composition (n=10-12 per group). I, Organ weight (n=15-17 per group). J, Images of iWAT and eWAT. Quantification of adipocytes surface areas (right, n=1025-1567 cells per group). (K-L) GTT (n=6-7 per group) and AUC (K), ITT (n=13-14 per group) and AUC (L) from mice on HFD for 2 weeks. Data are represented as mean ± s.e.m.; unpaired t-test or two-way ANOVA; * p<0.05, ** p<0.01, *** p<0.001.

We also crossed *Bckdk ^f/f^* mice with *Adipoq*-Cre transgenic mice to produce mature adipocyte-specific *Bckdk* knockout mice (*Adq*-*Bckdk*-AKO) (**Fig. S2O and S2P**). Compared with their littermates, HFD-fed *Adq*-*Bckdk*-AKO mice showed no significant difference in body weight gain, fat content, and organ mass (**Fig. S2Q-S2S**). Of note, the ratio of iWAT mass to body weight was increased in HFD-fed *Adq*-*Bckdk*-AKO mice (**Fig. 3B**).

**Fig. 3.**
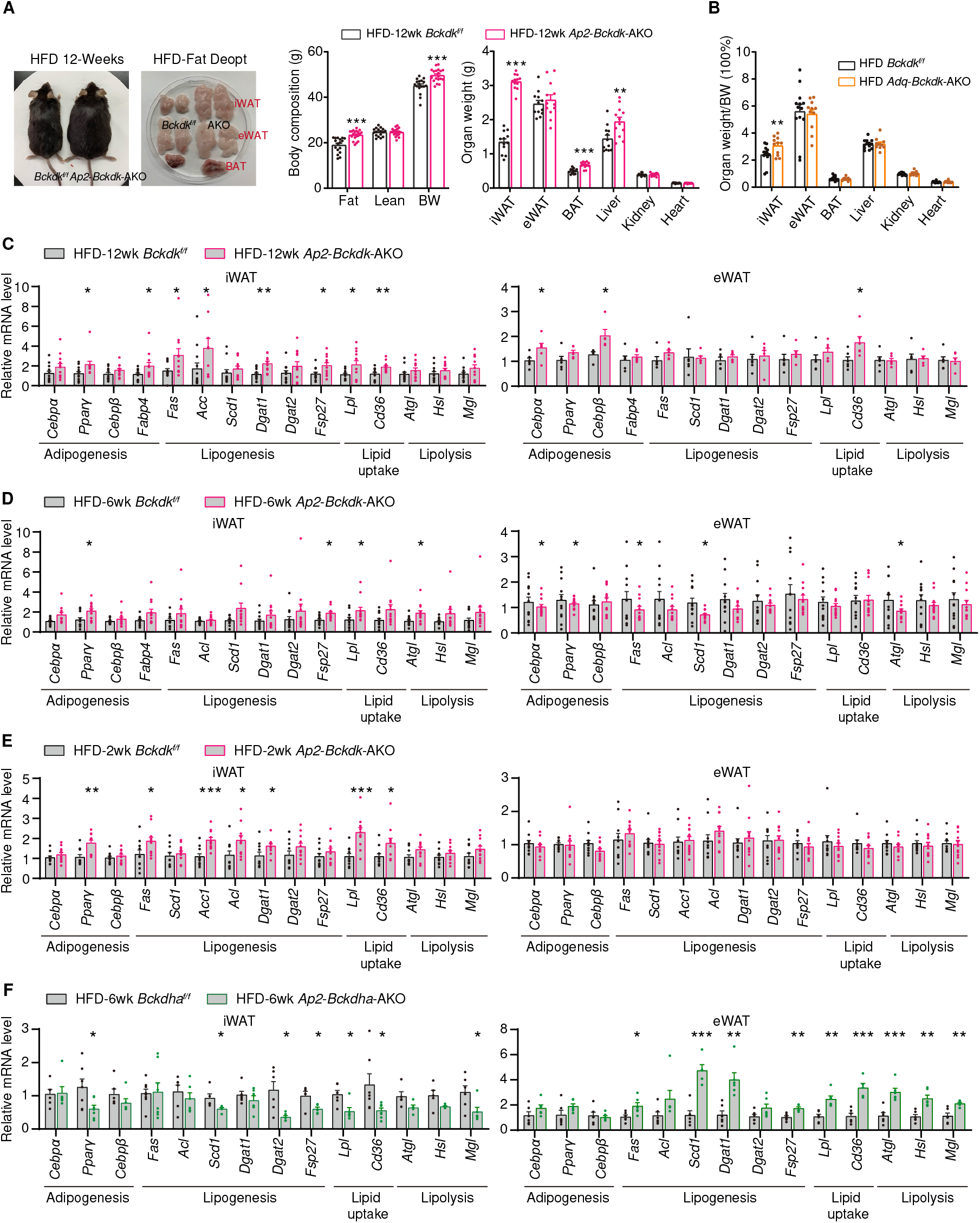
BCAA catabolism elicits depot-specific responses in sWAT and vWAT. (A) Representative photograph of mice and fat pads, body composition (n=17-19 per group), and organ weight (n=12 per group) of *Bckdk ^f/f^*and *Ap2*-*Bckdk*-AKO mice on HFD for 12 weeks. (B) Ratio of tissue mass to body weight in *Bckdk ^f/f^* and *Adq*-*Bckdk*-AKO mice on HFD for 6 weeks (n=11-13 per group). (C) mRNA expression of genes in the iWAT (n=12 per group) and eWAT (n=6 per group) from *Bckdk ^f/f^* and *Ap2*-*Bckdk*-AKO mice on HFD for 12 weeks. (D) mRNA expression of genes in the iWAT (n=11-12 per group) (A) and eWAT (n=9-12 per group) (B) from *Bckdk ^f/f^* and *Ap2*-*Bckdk*-AKO mice on HFD for 6 weeks. (E) qPCR analysis of the expression of indicated genes in the iWAT (n=10-11 per group) and eWAT (n=10-11 per group) of male *Bckdk^f/f^* and *Ap2-Bckdk*-AKO mice on high fat diet (HFD) for 2 weeks. (F) qPCR analysis of the expression of indicated genes in the iWAT (n=6-8 per group) and eWAT (n=5-6 per group) of male *Bckdha^f/f^* and *Ap2-Bckdha*-AKO mice on high fat diet (HFD) for 6 weeks. Data are represented as mean ± s.e.m.; unpaired t-test; * p<0.05, ** p<0.01, *** p<0.001.

### Blocking adipose BCAA catabolism attenuates diet-induced obesity in *Ap2*-*Bckdha*-AKO mice

We next analyzed the impacts of blocking BCAA catabolism on adipose physiology and obesity. We generated mice carrying the floxed *Bckdha* allele containing two loxP sites flanking exon 4 (*Bckdha ^f/f^*) and crossed them with *Fabp4*(*Ap2*)-Cre (**Fig. S3A**) transgenic mice. In *Ap2*-*Bckdha*-AKO mice, BCKDE1α expression was significantly reduced in iWAT, eWAT, and BAT, but not in other major tissues (**Fig. S3B**). On NCD, the weight gain, body composition, organ weights, glucose and insulin tolerance of *Ap2*-*Bckdha*-AKO mice were not different from those of control littermates (**Fig. S3C-S3G**). In response to HFD, however, the weight gain of *Ap2*-*Bckdha*-AKO mice was lower than that of control mice (**Fig. 2G**). *Ap2*-*Bckdha*-AKO mice demonstrated reduced fat but not lean mass (**Fig. 2H**), accompanied with lower mass of eWAT, iWAT, BAT, liver, but not kidney and heart (**Fig. 2I**), smaller adipocyte (**Fig. 2J**), better glucose clearance (**Fig. 2K**), and comparable insulin sensitivity (**Fig. 2L**), food intake, energy expenditure, and physical activity (**Fig. S3H-S3L**).

We also crossed *Bckdha^f/f^* mice with *Adipoq*-Cre transgenic mice to generate knockout mice with BCAA catabolic defect in mature adipocyte (*Adq*-*Bckdha*-AKO) (**Fig. S3M-3N**). Compared to control littermates consuming HFD, *Adq*-*Bckdha*-AKO mice did not show significant differences in adiposity (**Fig. S3O-3V**).

### Depot-specific impacts of BCAA catabolism on sWAT and vWAT

We noticed that, after 12-week HFD feeding, *Ap2*-*Bckdk*-AKO mice showed higher body fat content than control littermates, accompanied with significantly greater iWAT. Intriguingly, the eWAT mass showed no significant difference between the two mouse groups (**Fig. 3A**). On the other hand, compared with the littermates on HFD, *Adq*-*Bckdk*-AKO mice showed a modest increase in the ratio of iWAT mass to body weight while the eWAT was not affected (**Fig. 3B** and **Fig. S2Q-S2S**). These observations indicate that enhancing BCAA catabolism elicits fat-depot specific responses in iWAT and eWAT.

We next examined how BCAA catabolism affects the gene expression in these fat depots. In the iWAT of *Ap2-Bckdk*-AKO mice after 12-week HFD feeding, the expression of numerous lipogenic and adipogenic genes was significantly higher than that in control littermates, whereas few genes showed elevated expression in the eWAT (**Fig. 3C**). Furthermore, after 6-week HFD feeding, the expression of adipogenic and lipogenic genes was increased in the iWAT of *Ap2-Bckdk*-AKO mice compared with that in control littermates, but their expression in eWAT was unexpectedly reduced (**Fig. 3D**). After 2-week HFD feeding when the body weight and adipose mass showed no difference between *Ap2-Bckdk*-AKO and control mice, the expression of adipogenic and lipogenic genes was significantly higher in the iWAT but not eWAT in *Ap2-Bckdk*-AKO mice (**Fig. 3E**). On the other hand, in line with the attenuated obesity and fat mass in *Ap2-Bckdha*-AKO mice after 6 weeks of HFD feeding (**Fig. 2G-2I**), the expression of lipogenic and adipogenic genes was lower in the iWAT (**Fig. 3F**). In contrast, their expression was unexpectedly increased in the eWAT of *Ap2-Bckdha*-AKO mice (**Fig. 3F**). Together, these results demonstrate that BCAA catabolism elicits depot-specific responses in iWAT and eWAT. Of note, the adipogenic and lipogenic gene expression in iWAT but not eWAT is in line with the WAT expansion and development of obesity modulated by BCAA catabolism.

### BCAA catabolism promotes obesogenic adipogenesis in sWAT

WAT expansion in obesity occurs through mature adipocytes hypertrophy and/or precursors differentiation (adipogenesis) ^31–33^. *Adipoq*-cre specifically targets mature adipocytes while *Ap2*-Cre affects mature adipocytes and other cell types such as adipocyte precursors ^25, 28^. Thus, the different metabolic traits between *Ap2*-*Bckdk*-AKO and *Adq*-*Bckdk*-AKO mice or between *Ap2*-*Bckdha*-AKO and *Adq*-*Bckdha*-AKO mice (**Fig. 2** **and** **Fig. S2** **and** **S3**) indicated cell types other than mature adipocytes play a key role in the BCAA catabolism-regulated adipose physiology. A recent single cell sequencing study showed BCAA catabolic genes, along with the master regulators of adipogenesis, were among the hallmarker adipogenesis gene sets enriched in adipocyte progenitors ^34^. Given that the adipogenic regulators expression was higher in the iWAT of *Ap2-Bckdk*-AKO mice and lower in the iWAT of *Ap2-Bckdha*-AKO mice (**Fig. 3C-F**), we next investigated the impacts of BCAA catabolism on adipocyte progenitors and obesogenic adipogenesis in sWAT.

WAT composes of mature adipocytes and stromal-vascular fraction (SVF) cells, the latter include adipocyte precursor cells and various other cell types. *Bckdk* expression was significantly reduced in the SVF cells from the iWAT and eWAT of *Ap2*-*Bckdk*-AKO (**Fig. S4A and S4B)**. Importantly, the expression of adipogenesis markers and lipogenic genes was significantly elevated in SVF cells but not mature adipocytes from the iWAT of *Ap2*-*Bckdk*-AKO mice at the early stage of obesity, compared with those in control littermates (**Fig. 4A and 4B**). Meanwhile, no significant changes were observed in neither SVF cells nor mature adipocytes in the eWAT (**Fig. S4C and S4D**). On the other hand, in *Ap2*-*Bckdha*-AKO mice after 2 weeks of HFD feeding, the expression of *Pparγ* and *Cebpα* was lower in the SVF cells but not mature adipocytes from iWAT (**Fig. S4E and S4F**), and showed no changes in SVF cells and mature adipocytes from eWAT (**Fig. S4G and S4H**). The coordinated expression of adipogenic regulators in iWAT of *Ap2*-*Bckdk*-AKO and *Ap2*-*Bckdha*-AKO mice indicates BCAA catabolism enhances adipogenesis in sWAT.

**Fig. 4.**
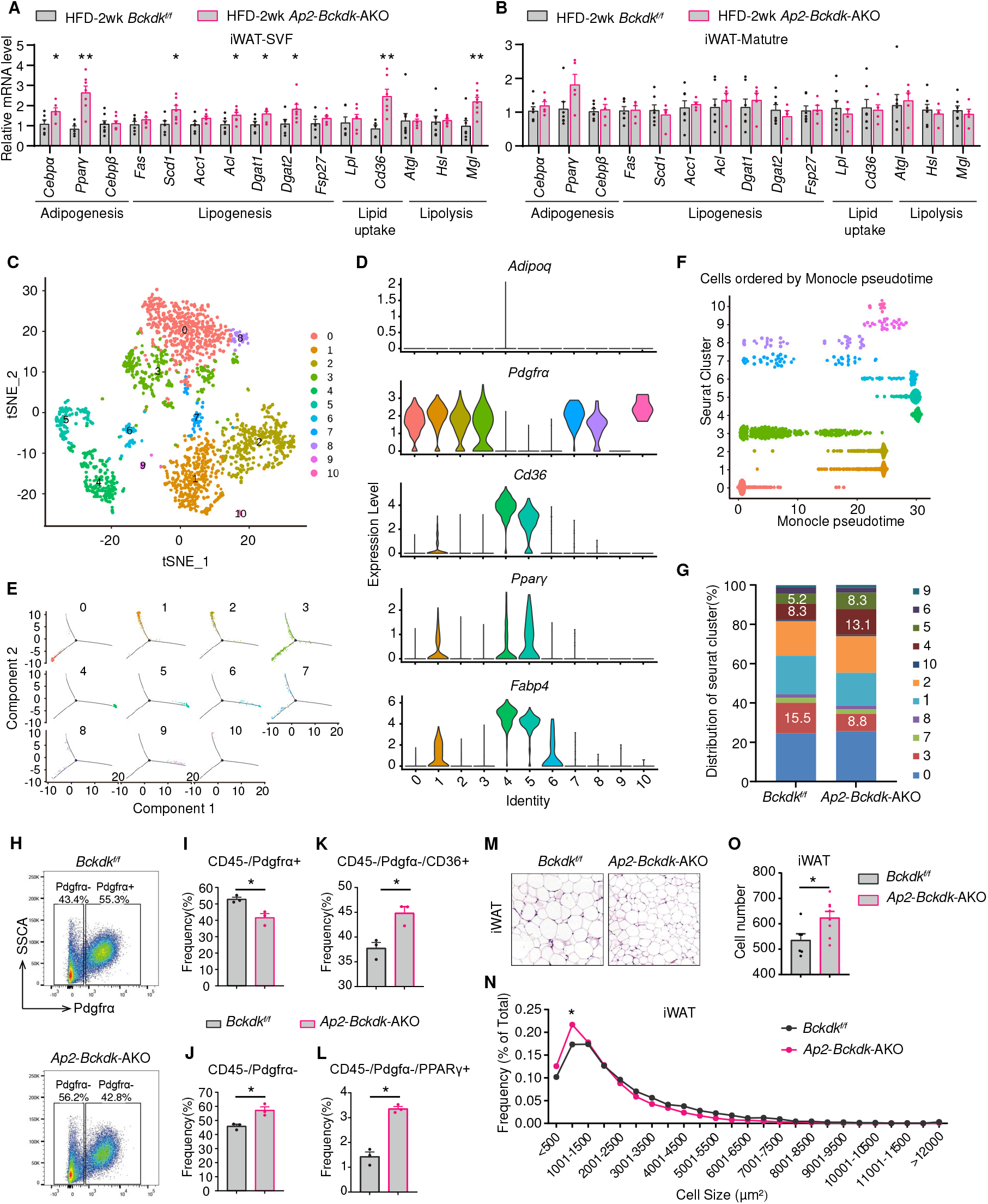
BCAA catabolism promotes obesogenic adipogenesis in sWAT. (A-B) mRNA expression of genes in the stromal vascular fraction (SVF) (A, n=5-8 per group) and mature adipocytes (Mature) (B, n=5-7 per group) of iWAT from *Bckdk ^f/f^* and *Ap2*-*Bckdk*-AKO mice on HFD for 2 weeks. (C) tSNE plot of combined CD45-negative SVF cells isolated from the iWAT of *Bckdk ^f/f^* and *Ap2*-*Bckdk*-AKO mice (n=3 per group) on HFD for 2 weeks. (D) Violin plots for markers. (E-F) Trajectory analysis (E) and pseudotime analysis (F) of combined CD45-negative SVF cells. (G) Group distribution of CD45-negative SVF cells from the iWAT of *Bckdk ^f/f^* and *Ap2*-*Bckdk*-AKO mice on HFD for 2 weeks. (H-L) Flow cytometry analysis of CD45-SVF cells in the iWAT of *Bckdk ^f/f^* and *Ap2*-*Bckdk*-AKO mice (n=3 per group) on HFD for 2 weeks. (M-O) Histology (M), adipocyte size (N), and adipocyte number (O) analysis in the iWAT of *Bckdk^f/f^* and *Ap2-Bckdk*-AKO mice (n=6-8 per group) on HFD for 2 weeks. Data are represented as mean ± s.e.m.; unpaired t-test; * p<0.05, ** p<0.01.

To further reveal the impact of BCAA catabolism on adipogenesis *in vivo*, we performed single-cell sequencing analysis of SVF cells from the iWAT of *Ap2*-*Bckdk*-AKO and control mice fed HFD for 2 weeks. Unsupervised clustering of the combined gene expression profiles identified 20 clusters (**Fig. S4I and S4J**). Of note, most of these clusters, with the exception of Cluster 3, 4, 8, and 20, were marked by high expression of leukocytes marker CD45 (encoded by *Ptprc*) (**Fig. S4K and S4L**). Based on the expression of canonical markers, the top three abundant clusters are B cells (Cluster 0, marked by *Cd19* and *Ms4a1*), CD4 T cells (Cluster 1), and CD8 T cells (Cluster 2), respectively (**Fig. S4M**).

Next, we re-analyzed the CD45-SVF cells, i.e., the Cluster 3, 4, 8, and 20 cells aforementioned (**Fig. S4N**). Unsupervised clustering of the combined gene expression profiles of these cells identified 10 groups (G1–10) (**Fig. 4C** **and S4O**). G0, G1, G2, G3, G7, G8 and G10 cells were marked by high level expression of adipose progenitor markers *Pdgfrα* (**Fig. 4D**). Among the *Pdgfrα*-groups, G4 and G5 cells were marked by adipocyte identity genes including *Pparγ*, *Fabp4*, and *Cd36*. A weak expression of adiponectin was detected in G4 (**Fig. 4D**). The pseudotemporal analysis indicated that G0 cells provide a source for G3 cells, which have two differentiation trajectories for either G1 and G2 cells or G4 and G5 cells, respectively (**Fig. 4E-4F** **and S4P**). Further analysis of the group distribution of CD45-SVF cells in *Ap2*-*Bckdk*-AKO and control mice showed the G3 proportion was decreased from 15.5% (control) to 8.8% (*Ap2*-*Bckdk*-AKO). In contrast, the proportions of G4 and G5 cells were increased from 8.3% and 5.2% (control) to 13.1% and 8.3% (*Ap2*-*Bckdk*-AKO), respectively (**Fig. 4G**). The proportions of other groups remained similar. The population shifts indicate more G3 cells differentiate into G4 and G5 cells in the sWAT of *Ap2*-*Bckdk*-AKO.

To further characterize the CD45-SVF cell subpopulations in the sWAT, we performed flow cytometry analysis using antibodies against PDGFRα, PPARγ, and CD36. Compared with that in control mice, the proportion of PDGFRα+ cell was reduced while the proportion of PDGFRα-cells was increased in the sWAT of *Ap2*-*Bckdk*-AKO mice after 2 weeks of HFD feeding (**Fig. 4H-4J**), accompanied with significantly increased proportions of PDGFRα-/PPARγ+ and PDGFRα-/CD36+ cells (**Fig. 4K-4L**). These results support enhanced obesogenic adipogenesis in the sWAT of *Ap2*-*Bckdk*-AKO mice.

Next, histological analysis demonstrated greater adipocyte number and higher portion of small adipocytes in the iWAT of *Ap2*-*Bckdk*-AKO mice after 2-week-HFD-feeding, compared with those in control mice (**Fig. 4M-4O**), while the changes in eWAT were not significant (**Fig. S4Q-S4S**), further suggesting adipogenesis in sWAT is enhanced by BCAA catabolism. This is intriguing as it has been reported that HFD does not induce adipocyte hyperplasia in the sWAT ^35, 36^.

### An intermediate of leucine catabolism governs the adipogenic transcriptional cascade

In adipose tissue, in addition to mature adipocytes and their progenitors, *Fabp4*-Cre activity has also been detected in other cell types such as endothelial cells and macrophages. We then assessed whether BCAA catabolism affected adipocyte progenitor differentiation in a cell-autonomous fashion by isolating SVF cells from the iWAT of *Ap2*-*Bckdk*-AKO and *Ap2*-*Bckdha*-AKO mice and differentiating them into mature adipocytes *in vitro*. As expected, *Bckdk* deficiency enhanced, while *Bckdha* deficiency suppressed, PPARγ expression and adipocyte differentiation (**Fig. 5A-5F**).

**Fig. 5.**
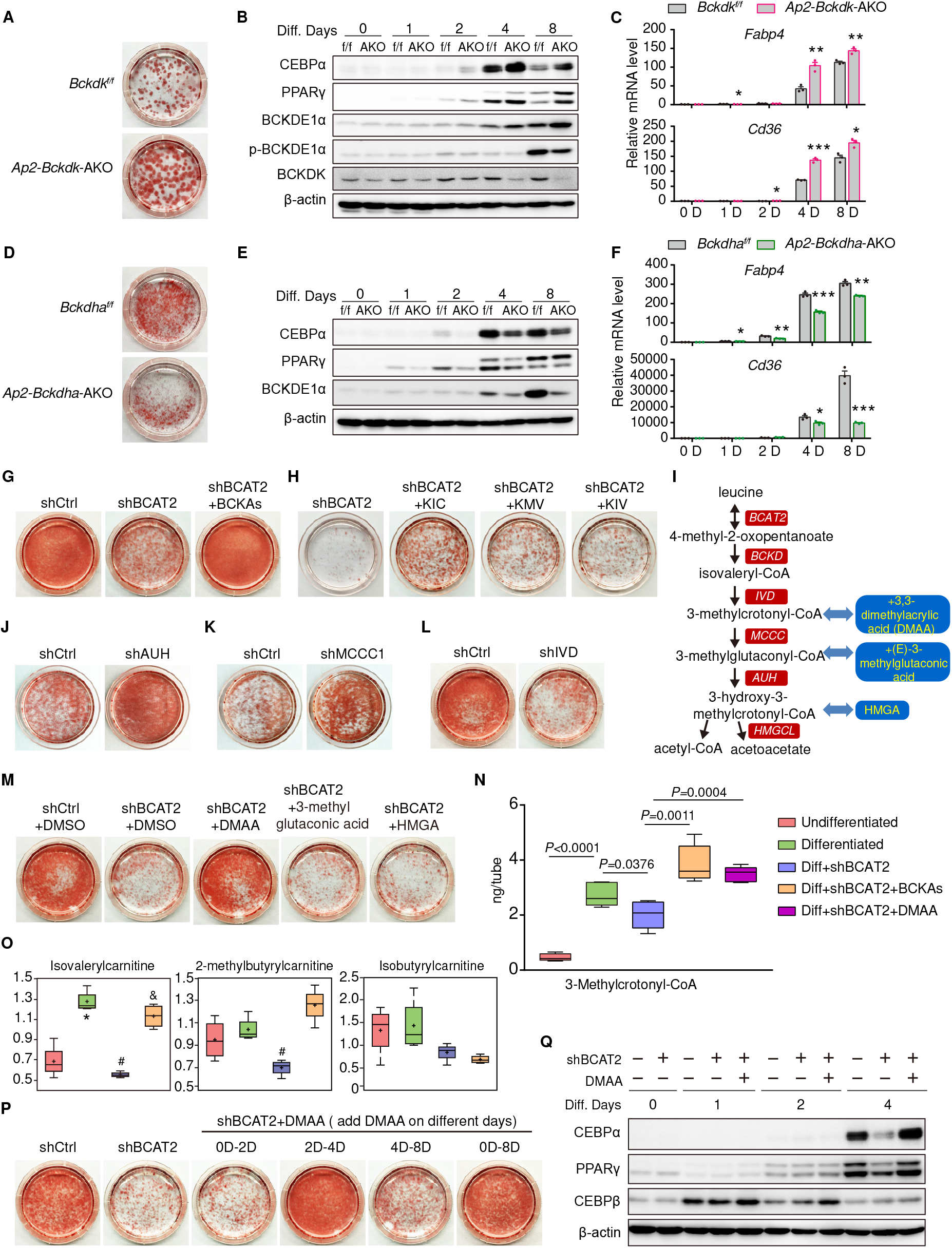
A leucine catabolic intermediate governs adipocyte differentiation. (A) Representative images of oil red staining of differentiated SVF cells from iWAT of *Bckdk ^f/f^* and *Ap2*-*Bckdk*-AKO mice. (B-C) Protein and mRNA expression during SVF cell differentiation. (D) Representative images of oil red staining of differentiated SVF cells from *Bckdha ^f/f^* and *Ap2*-*Bckdha*-AKO mice. (E-F) Protein and mRNA expression during SVF cell differentiation. (G-H) Representative images of oil red staining of differentiated 3T3-L1 adipocytes with *Bcat2* silencing in the presence or absence of three BCKAs (500 μM each) (G) or individual BCKA (500 μM) (H) supplementation. (I) Leucine catabolic process. (J-M) Representative images of oil red staining of differentiated 3T3-L1 adipocytes with: shRNA silencing of *Auh* (J), *Mccc*1 (K), *Ivd* (L), *Bcat2* silencing with or without different leucine catabolite derivatives (M). (N) Abundances of leucine catabolite 3-methylcrotonyl-CoA in 3T3-L1 adipocytes on the 4th day of differentiation. (O) Metabolomics analysis of BCAA catabolites: isovaleryl-carnitine from leucine, 2-methylbutyryl-carnitine from isoleucine, and isobutyryl-carnitine from valine in 3T3-L1 adipocytes on the 4th day of differentiation. *p<0.05 vs Undifferentiated group; #p<0.05 vs Differentiated group; &p<0.05 vs Differentiated+shBCAT2 group. (P) Representative images of oil red staining of differentiated 3T3-L1 adipocytes with *Bcat2* silencing, treated with DMAA (5 mM) during different periods of differentiation. (Q) Protein expression during 3T3-L1 differentiation. Data are represented as mean ± s.e.m.; unpaired t-test or one-way ANOVA; * p<0.05, ** p<0.01, *** p<0.001.

3T3-L1 adipocyte differentiation model recapitulates many of the key adipogenic and lipogenic processes occurring *in vivo*. We found that, among all proteogenic amino acids, BCAAs demonstrated uniformly the highest rate of consumption starting from the intermediate differentiation stage (**Fig. S5A-S5B**). It has been reported that BCAA catabolism is essential for 3T3-L1 differentiation ^22, 24^. Indeed, blocking BCAA catabolism by inactivating BCAT2 significantly suppressed 3T3-L1 differentiation and supplementing BCKAs to the *Bcat2*-silenced 3T3-L1 cells resumed BCAA catabolism and adipocyte differentiation (**Fig. 5G** **and S5C-S5E**). The replenishment of leucine keto acid metabolite (KIC) demonstrated the strongest restoration of adipocyte differentiation in *Bcat2*-silenced 3T3-L1 cells, compared to the keto acid metabolites of isoleucine (KMV) and valine (KIV) (**Fig. 5H**). These data validated the essential role of BCAA, particularly leucine, catabolism in adipocyte differentiation.

Reports suggested leucine-derived acetyl-CoA provided substrates for fatty acid synthesis during adipocyte differentiation (**Fig. 5I**) ^17, 18^. Surprisingly, our results showed that blocking the acetyl-CoA production from leucine by silencing the distal downstream enzyme MCCC or AUH resulted in enhanced rather than inhibited adipocyte differentiation (**Fig. 5J**-**5K** **and S5F-S5G**). Meanwhile, inactivating the upstream enzyme IVD (**Fig. 5L** **and S5H**) potently inhibited adipocyte differentiation. These results suggested that the IVD-produced metabolic intermediate of leucine played a critical role in promoting adipocyte differentiation. In 3T3-L1 cells with *Bcat2* silenced, the administration of +3,3-dimethylacrylic acid (named DMAA, a derivative of IVD product 3-methylcrotonyl-CoA) showed the strongest restoration of adipocyte differentiation, compared to MCCC or AUH products (**Fig. 5M**). These results support a potent and unique differentiation-promoting function of 3-methylcrotonyl-CoA or its derivatives.

During 3T3-L1 cell differentiation, the abundances of 3-methylcrotonyl-CoA and isovalerylcarnitine generated from leucine catabolism were dramatically elevated, which was abolished by *Bcat2* inactivation and restored by BCKA or DMAA replenishment (**Fig. 5N and 5O**). Importantly, the changes in the levels of leucine catabolites were in agreement with the differentiation (**Fig. 5G and 5M**). The elevation of leucine catabolites in differentiating adipocyte was accompanied with high level expression of the upstream enzymatic genes (*Bcat2*, *Ppm1k, Bckdha, Bckdhb, Dbt,* and *Ivd*) but minimally-induced expression of the distal downstream enzymatic genes (*Auh* and *Mccc1*) in the leucine catabolic pathway (**Fig. 5I** **and S5I-S5K**).

3T3-L1 adipocyte differentiation involves several defined stages and a coordinated transcriptional cascade ^37, 38^. In *Bcat2*-silenced 3T3-L1 cells, the rescuing effects of DMAA or BCKA were tested during different stages, and found to be effective during the Day 2 to Day 4 period of differentiation (**Fig. 5P** **and S5L**). This period is the critical stage of cell fate determination when the master regulators of adipogenesis such as C/EBPα and PPARγ are induced ^39^. Indeed, inactivation of *Bcat2* abolished the expression of C/EBPα and PPARγ, which was restored by DMAA or BCKA supplementation at the intermediate differentiation stage (**Fig. 5Q** **and S5M-S5N**). BCAA catabolism did not affect mitotic clonal expansion (**Fig. S5O**) and the expression of other differentiation regulators including C/EBPβ or *Chop* (**Fig. S5M-S5N and S5P**). Therefore, a leucine intermediate metabolite plays a regulatory role in adipocyte differentiation via governing the expression of adipogenic master regulators and the transcriptional cascade.

### Leucine catabolism promotes adipocyte differentiation via polyamine

To better understand how BCAA catabolism regulated adipocyte differentiation, we performed metabolomics analyses in differentiating 3T3-L1 cells in which BCAA catabolism was blocked or rescued (**Fig. S6A-S6C**). Interestingly, among >200 metabolites involving in carbohydrate, lipid, amino acids, and nuclear acids metabolism, only polyamines showed striking changes associated with BCAA catabolism (**Fig. 6A**). Polyamines, including putrescine, spermidine, and spermine, demonstrated significantly elevated abundances in differentiating adipocytes, which were abolished by *Bcat2* silencing but restored by the supplement of BCKAs (**Fig. 6A**). The changes of polyamine abundances were further supported by the changes in the protein levels of spermidine/spermine N1-acetyltransferase (SSAT) (**Fig. 6B** **and S6D**). SSAT is the rate-limiting enzyme of polyamine degradation and its protein expression is induced by polyamines ^40^.

**Fig. 6.**
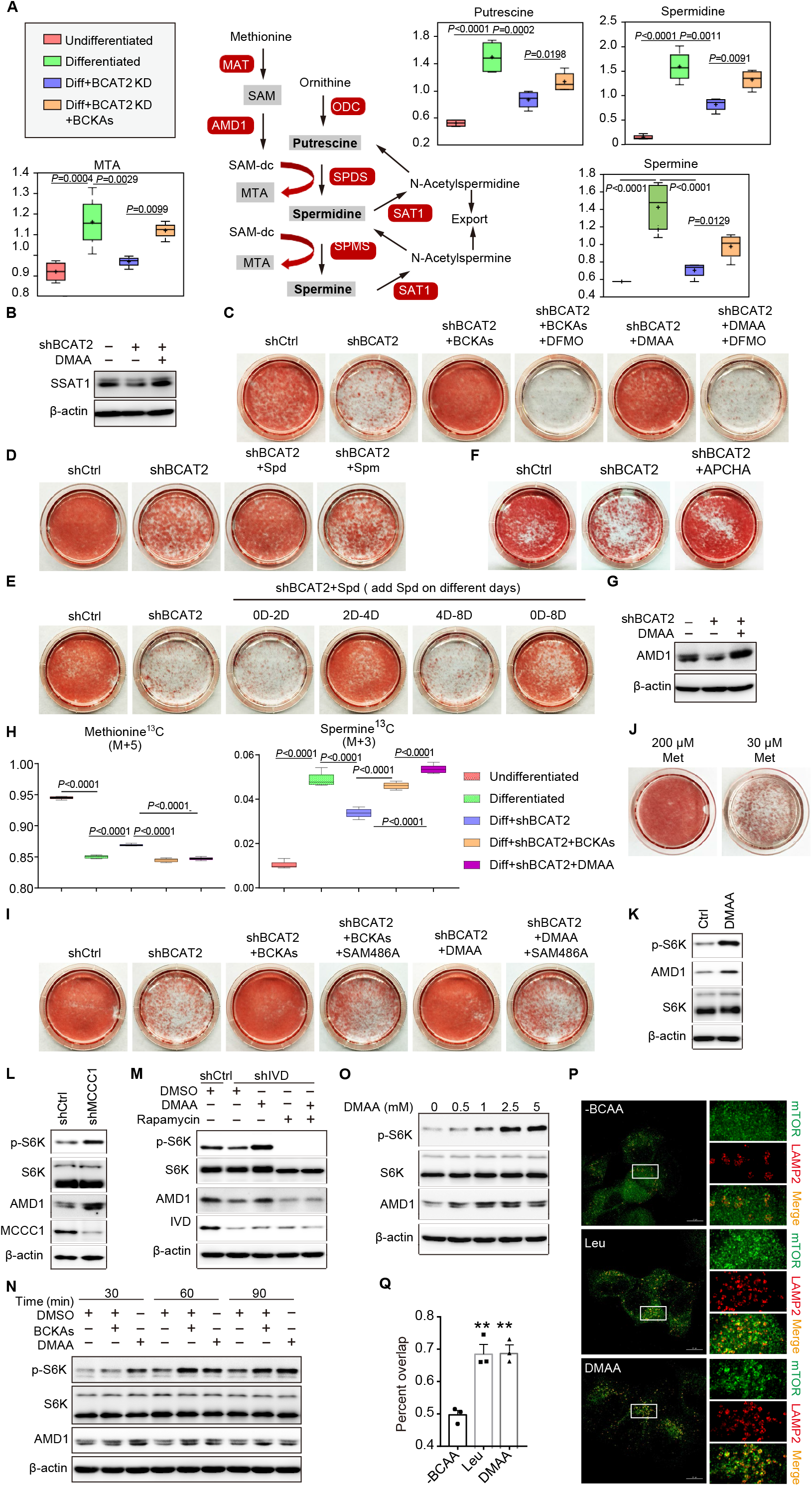
The leucine catabolic intermediate activates mTORC1 and methionine-AMD1-driven polyamine synthesis to promotes adipocyte differentiation. (A) Metabolomics analysis of polyamines in undifferentiated and differentiating 3T3-L1 adipocytes with *Bcat2* silencing or BCKAs treatment (n=4 per group) on the 4th day of differentiation. (B) SSAT1 protein expression. (C-F) Representative images of oil red staining of differentiated 3T3-L1 adipocytes with *Bcat2* silencing, treated with BCKAs (C), DMAA (C), DFMO (50 μM, C), spermine (Spm) (7 μM, D), spermidine (Spd) (10 μM, D), Spd during different periods (E), APCHA (100 μM, F). (G) AMD1 protein expression. (H) Measurement of polyamine synthesis using L-Methionine-^13^C5,^15^N. The y-axis represents the relative mass isotope distribution (%). (I-J) Representative images of oil red staining of differentiated 3T3-L1 adipocytes treated with SAM486A (10 μM), BCKAs, and DMAA (I), low methionine in medium (30 μM, J). (K-M) Western blotting of AMD1 and mTORC1 signaling in differentiating 3T3-L1 adipocytes with DMAA treatment (K), *Mccc1* silencing (L), *Ivd* silencing and DMAA and rapamycin (20 nM, 6 hour) treatment (M). (N-O) AMD1 protein and mTOR signaling in undifferentiated 3T3-L1 pre-adipocytes treated at different time points (N) or various concentration for 1 hour (O) in BCAA-free medium. (P-Q) Representative images of mTOR and LAMP2 localization in Hela cells treated with leucine or DMAA in BCAA-free medium (P), with quantification (Q, 20 cells). Data are represented as mean ± s.e.m.; one-way ANOVA; ** p<0.01.

Previous studies have demonstrated that polyamine, particularly spermidine, governs C/EBPα and PPARγ expression and adipocyte differentiation ^41–44^. To investigate the role of polyamine in BCAA catabolism-regulated adipocyte differentiation, α-difluoromethylornithine (DFMO), an inhibitor of the rate-limiting enzyme for polyamine biosynthesis ornithine decarboxylase (ODC), was used to block polyamine production. DFMO treatment markedly abolished BCKA-or DMAA-dependent adipocyte differentiation in the *Bcat2*-silenced 3T3-L1 cells (**Fig. 6C**). On the other hand, while the blockade of BCAA catabolism diminished adipocyte differentiation and polyamine abundances (**Fig. 6A**), spermidine replenishment restored C/EBPα and PPARγ expression and the differentiation of *Bcat2*-inactivated 3T3-L1 cells (**Fig. 6D** **and S6E**). Consistent with BCKA and DMAA supplement, treatment of spermidine during Day 2 to Day 4 restored *Bcat2*-silencing-impaired 3T3-L1 differentiation and C/EBPα and PPARγ expression (**Fig. 6E** **and S6F**). Furthermore, N-(3-aminopropyl)-cyclohexylamine (APCHA), a chemical inhibitor of spermine synthase, has been reported to enhance adipocyte differentiation by increasing the ratio of spermidine/spermine ^43^. Indeed, APCHA treatment also restored differentiation in the *Bcat2*-silenced 3T3-L1 cells (**Fig. 6F**). Thus, polyamine, particularly spermidine, mediates the DMAA’s effect on promoting C/EBPα and PPARγ expression and adipocyte differentiation.

### DMAA promotes polyamine synthesis from methionine

We next investigated how DMAA increased polyamine abundance. Polyamines are synthesized from methionine and ornithine with two rate-limiting enzymes, S-adenosylmethionine decarboxylases (AMD) and ODC, respectively (**Fig. 6A**). BCAA catabolic manipulations or DMAA treatment failed to affect the expression of ODC in differentiating 3T3-L1 adipocytes (**Fig. S6G-S6H**). AMD1 protein level was reduced by *Bcat2* silencing, which was restored by DMAA or BCKA supplementation (**Fig. 6G** **and S6I**), suggesting that leucine catabolism promoted AMD1 expression. This relationship was also supported by the BCAA-catabolism-dependent fluctuation of methylthioadenosine (MTA), a by-product of polyamine synthesis from methionine (**Fig. 6A**). We used radiolabeled L-Methionine-^13^C5,^15^N to quantify the polyamine synthesis. The results showed that spermine was synthesized from methionine, which was enhanced in differentiating adipocytes. Importantly, blocking BCAA catabolism inhibited polyamine synthesis from methionine, which was rescued by DMAA or BCKA supplementation in culture medium (**Fig. 6H**). Furthermore, inhibition of AMD1 with SAM486A ^45^ abolished BCKAs-or DMAA-dependent adipocyte differentiation (**Fig. 6I**), suggesting AMD-mediated polyamine synthesis from methionine was essential for DMAA-promoted differentiation. Indeed, in addition to BCAAs, methionine was another essential amino acid showing substantial consumption during adipocyte differentiation (**Fig. S5A**). Limiting methionine supply in culture medium significantly impaired adipocyte differentiation (**Fig. 6J**). The expression of other polyamine synthetic enzymes was not affected by BCAA catabolism in 3T3-L1 adipocytes (**Fig. S6H**). Ornithine supplementation failed to rescue the impaired differentiation of *Bcat2*-silenced 3T3-L1 cells (**Fig. S6J**). Together, these results show that DMAA promotes polyamine synthesis from methionine to enhance adipocyte differentiation.

### DMAA activates mTORC1 to promote AMD1 expression

A recent report showed AMD1 protein was stabilized by the mechanistic target of rapamycin complex 1 (mTORC1) ^45^. In differentiating adipocytes, DMAA treatment markedly increased mTORC1 activity and AMD1 protein expression (**Fig. 6K**). Inactivation of MCCC1, the degrading enzyme for 3-methylcrotonyl-CoA, also increased mTORC1 activity and AMD1 protein expression (**Fig. 6L**). Blocking leucine catabolism via *Ivd*-or *Bcat2*-silencing resulted in lower mTORC1 activity and AMD1 protein level, which were restored by DMAA treatment. Importantly, mTORC1 inhibitor rapamycin abolished DMAA-induced AMD1 protein expression (**Fig. 6M** **and S6K**). In undifferentiated 3T3L-1 cells, BCKAs and DMAA were capable of stimulating mTORC1 quickly in the absence of BCAAs in culture medium (**Fig. 6N**). DMAA promoted AMD1 expression and mTORC1 activity in a dose-dependent way (**Fig. 6O**). Interestingly, AKT activity was suppressed by BCKAs and DMAA (**Fig. S6L**), indicating an AKT-independent mTORC1 activation by DMAA. DMAA treatment induced the recruitment of mTORC1 to the surface of lysosomes (**Fig. 6P and 6Q**), a key process for mTORC1 activation by leucine ^46^. The mechanism underlying DMAA-induced mTORC1 activation warrants further investigation.

## DISCUSSION

The current study demonstrates a regulatory role of BCAA catabolism in adipogenesis and WAT physiology. The unbiased genetic analyses reveal a link between the adipose BCAA catabolic pathway and obesity traits in mice and humans. While BCAA catabolism in mature adipocytes exerts minor effects on adiposity, enhancing adipose BCAA catabolism via FABP4-Cre-mediated *Bckdk* deletion promotes diet-induced obesity and WAT expansion in mice. Of interest, the sWAT responds to BCAA catabolism differently from eWAT. Adipogenesis in sWAT is promoted by BCAA catabolism in a cell-autonomous manner. Furthermore, the pro-adipogenic effect of BCAA catabolism is mediated by a leucine catabolic intermediate, but not BCAA *per se* nor the end product acetyl-CoA. Mechanistically, the catabolic intermediate, named DMAA, activates mTORC1 signaling and AMD1-methionine-driven polyamine synthesis to promote PPARγ expression and adipocyte differentiation (**Fig. S6M**).

The obesity-promoting function of BCAA catabolism in *Ap2*-*Bckdk*-AKO mice is intriguing. The reduced energy expenditure may contribute to the promoted obesity. A recent report showing blocking adipose BCAA catabolism via genetic deletion of *Bcat2* protects mice from diet-induced obesity by increasing sWAT browning and thermogenesis ^21^. However, in *Adq*-*Bckdha*-AKO mice, blocking adipose BCAA catabolism via *Bckdha* deletion shows no effect on diet-induced obesity. Given that both *Bcat2* and *Bckdha* deletion leads to BCAA catabolic defect in mature adipocytes in these mouse models, the cause for these discrepancies remains elusive. On the other hand, the BCAA-catabolism-promoted adipogenesis may contribute, at least in part, to the accelerated obesity in *Ap2*-*Bckdk*-AKO mice. Together, these results indicate that BCAA nutrient and their catabolism may accelerate obesity in response to excess caloric intake. This hypothesis is supported by a recent study showing that restriction of BCAA prevents diet-induced obesity in mice ^47^.

WAT plays a beneficial role in physiological conditions, which can disappear and become detrimental in pathological obesity due to inflammation, ectopic lipid deposition, amongst others. It has been repeatedly observed that BCAA catabolism is suppressed in WAT in obese animals and humans. Our current data show that, during the development of obesity, BCAA catabolism accelerates tissue expansion and lipid storage in WAT. It is plausible that the adipose BCAA catabolic defect in obese animals impedes further lipid storage in WAT and contributes to the ectopic lipid deposition. Meanwhile, the impaired BCAA catabolism in adipose tissue contributes to the elevated plasma BCAA. As ectopic lipid deposition and rising circulating BCAA may represent two major causes of insulin resistance, our finding here underscores a potential mechanism to the strong linkage observed between disrupted BCAA homeostasis and obesity-associated diseases.

BAT is specialized for energy expenditure. A recent study shows that BCAA catabolic defect in BAT attenuates fuel oxidation and thermogenesis, leading to higher weight gain and worsened glucose tolerance upon chronic metabolic challenge ^19^. However, the disruption of BCAA catabolism in both WAT and BAT in the *Ap2*-*Bckdha*-AKO mice results in less body weight gain and improved glucose tolerance. Thus, the functional changes of WAT, but not BAT, likely accounts for the reduced obesity in *Ap2*-*Bckdha*-AKO mice. Likewise, the enhanced BCAA catabolism in WAT likely determines the exacerbated obesity traits in *Ap2*-*Bckdk*-AKO mice. Meanwhile, BCAA catabolism reinforces energy expenditure in BAT and energy storage in WAT, which could counteract each other’s influences on systemic metabolism and contribute to the BCAA’s paradoxical impacts.

To make it more interesting, even for different white fat depots, BCAA catabolism exerts discrete effects on tissue expansion, adipogenesis, and lipid metabolic gene expression in sWAT and vWAT. The depot-specific impacts are intriguing given the different roles of these depots in health and pathology. Further investigation is warranted to find out how BCAA catabolism affects the inherent properties of these WAT depots and/or the crosstalk between them. In addition, while sWAT expansion is often considered be metabolically healthy by improving insulin sensitivity, the enhanced vWAT expansion may help to explain the enhanced insulin resistance in *Ap2*-*Bckdk*-AKO mice. Whether the further expansion of sWAT but not eWAT in the late stage of obesity results in metabolically healthy *Ap2*-*Bckdk*-AKO mice remains to be determined.

It has been suggested that the sWAT expands through adipocyte hypertrophy, while the vWAT expands by both hypertrophy and hyperplasia in response to HFD feeding in mice ^35, 36^. However, another report suggests vWAT grows mostly by hypertrophy and sWAT by hyperplasia ^48^. Meanwhile, numerous reports suggest that adipocyte progenitors derived from sWAT have higher adipogenic potential than those from vWAT, while some studies report the opposite or no difference ^49^. Thus, there is currently no clear consensus regarding the differences in the intrinsic adipogenic potential between depot-specific adipocyte progenitor cells. Our data suggest enhancing BCAA catabolism promotes the adipogenesis in sWAT, but not vWAT, in response to HFD feeding. sWAT and vWAT depots are intrinsically different, showing distinct cellularity, gene expression pattern, lipolytic rate, insulin sensitivity, and vascularization. It has been shown that the depot-specific microenvironment may also play a role in regulating adipogenesis ^50^. Further studies are warranted to clarify the interplay between the BCAA catabolism and the extrinsic factors in different fat depots during the development of obesity. Nevertheless, it is intriguing that BCKDK and the associated low BCAA catabolic activity may act as the gatekeeper for adipogenesis in sWAT.

The impact of BCAA catabolism on adipocyte differentiation leads to the obvious question about the underlying molecular basis. It has been shown that BCAA catabolism provides the sources of priming substrates for *de novel* synthesis of fatty acids in differentiating adipocytes *in vitro* ^17, 18^. In the present study, genetic inhibition of MCCC and AUH to block the leucine converting into acetyl-CoA enhances rather than suppresses adipocyte differentiation. In addition, DMAA, but not the MCCC and AUH products, promotes adipocyte differentiation and C/EBPα and PPARγ expression on 2-4 days of 3T3-L1 differentiation before the lipid accumulation stage. These results suggest that, while BCAA catabolism contributes to the lipogenic acetyl-CoA pool, the leucine catabolic intermediate produced by IVD is a potent regulator of adipocyte differentiation. Thus, the BCAA intermediate catabolites, such as DMAA and HIB, serve diverse regulatory functions in different metabolic processes.

Although simple in structure, polyamines are essential molecules for growth in eukaryotic cells and key regulators of adipocyte differentiation and metabolism. Our data show that spermidine mediates the effect of BCAA catabolism on PPARγ and C/EBPα expression and adipocyte differentiation. The leucine catabolic intermediate drives polyamine synthesis through the AMD-methionine branch. Indeed, in addition to BCAAs, methionine is another amino acid showing strong impacts on glucose and lipid metabolism ^51–53^. The cross-talks among BCAA, methionine, and polyamine metabolism are intriguing given their extensive metabolic impacts. These interplays highlight shared mechanisms.

mTORC1 activation by DMAA is of interest considering both leucine *per se* and leucine-derived acetyl-CoA and monomethyl branched-chain fatty acid are mTORC1 stimulators ^16, 54–56^. DMAA quickly activates mTORC1 in the absence of BCAA in culture medium. Similar to leucine, DMAA induces the localization of mTORC1 to the surface of lysosomes. It remains unclear whether DMAA *per se* activates mTORC1 or it needs to be converted to other metabolites such as 3-methylcrotonyl-CoA. Nevertheless, it is intriguing that both leucine and its catabolites can activate mTORC1. How these signals act coordinately with different kinetics, in different tissues, under different physiological and pathological conditions warrants further investigation.

In summary, the current study demonstrates a previously unknown role of BCAA catabolism as a regulator of adipogenesis, WAT physiology, and the development of obesity. The intricate functions of BCAA catabolism in different adipose depots may exert diverse effects on the system metabolism, highlighting the temporal and spatial complexity of understanding the BCAA’s controversial metabolic impacts. Moreover, the DMAA-driven signaling and metabolic network underlying leucine, methionine, and polyamine metabolism sheds lights on the regulatory function of bioactive intermediate metabolites and the coordinated cross-talk among amino acids in the development of obesity. In addition to amino acids *per se*, much attention may be paid on their tissue- and cell-specific metabolic fate to gain a comprehensive understanding of their roles in metabolic health and diseases.

## MATERIALS and METHODS

### Gene-trait correlation in Hybrid Mouse Diversity Panel (HMDP)

We accessed the correlation data between genes and metabolic traits measured in HMDP (https://systems.genetics.ucla.edu/data/hmdp) comprised of ∼100 mouse strains fed with chow or high-fat diet ^57, 58^. Gene-trait correlation for a total of ten metabolic traits were collected, including body weight, fat mass, adiposity, gonadal fat, mesenteric fat, retroperitoneal fat, LDL, HDL, total cholesterol and triglyceride. Metabolic traits were assessed using 8-12 mice per strain, whereas adipose tissue expression profiling was performed using 3 mice per strain.

### Human GWAS for lipid metabolism

We collected publicly available human GWAS data for triglyceride (TG) and body mass index (BMI) from large meta-analysis consortia including the Global Lipids and Genetics Consortium (GLGC) ^59^ and the Genetic Investigation of Anthropometric Traits consortium (GIANT) ^60^. Summary level statistics for all single nucleotide polymorphisms (SNPs) analyzed were obtained from each study. For each human GWAS dataset, the summary-level statistics for all SNPs was filtered by removing the weakly associated SNPs (lower 80%) and SNPs in high linkage disequilibrium (r2>0.5).

### Adipose coexpression modules

A total of 491 adipose coexpression modules (ACMs) were constructed from multiple human and mouse studies (Supplementary Table 2) using WGCNA ^61^. For each ACM, functional enrichment of canonical pathways (described below) was assessed using Fisher’s exact test. Pathways reaching Bonferroni corrected p < 0.05 was determined to be significant.

### Canonical pathways

Canonical pathways, including twelve amino acid metabolic pathways, were retrieved from Kyoto Encyclopedia of genes and genomes (KEGG) ^62^ and Reactome ^63^. We manually added *PPM1K* and *BCKDK* into the BCAA pathway due to their key regulatory role in BCAA catabolism. BCAA pathway was further categorized into genes specific to degradation of leucine, valine and isoleucine, respectively, yielding a total of 15 amino acid pathways.

### Adipose expression quantitative trait loci (eQTLs)

We curated a comprehensive list of human adipose tissues (subcutaneous and visceral) eQTLs from individual studies as well as large consortia such as the Genotype-Tissue Expression (GTEx) project ^64^ and the Multiple Tissue Human Expression Resource (MuTHER) consortium ^65^. All eQTL studies included are listed in Supplementary Table 2.

### Adipose Bayesian networks

Adipose Bayesian networks were constructed using genetic and expression profiling data from adipose tissues of both human and mouse samples (Supplementary Table 2) through a previously developed method ^66, 67^. We further merged our Bayesian networks with the adipose functional interaction networks constructed from thousands of diverse experiments ^68^.

### Marker set enrichment analysis

We used the marker set enrichment analysis (MSEA) module provided in the Mergeomics pipeline ^69^ to determine the genetic association strength of the ACMs and individual amino acid pathways with human metabolic traits. The ACMs and amino acid pathways were first mapped to adipose eQTLs to derive the corresponding representative eSNP sets. The disease association p values of the adipose eSNPs representing each module or pathway were then extracted from each human GWAS study and tested for significant enrichment of disease-associated eSNPs using a chi-square like statistics. False discovery rate (FDR) was determined based permutation and FDR < 15% was used as the significance cutoff. Analysis was performed for subcutaneous adipose eQTLs, visceral adipose eQTLs and combined adipose eQTLs separately, and the minimum FDR out of the three sets of analyses was used to represent the significance of each ACM.

### Key Driver Analysis

The weighted Key Driver Analysis (KDA) module in the Mergeomics pipeline ^69^ was applied to pinpoint key drivers of BCAA ACMs. Key driver (KD) is defined as the gene whose neighboring subnetwork exhibited significant enrichment for ACM genes, and KD is more likely to play important regulatory role on ACM. KDA firstly maps ACM genes to weighted adipose tissue Bayesian networks, then utilized a chi-square like statistics to test for enrichment of ACM genes, taking the edge weight information (consistency score across networks from different datasets) into consideration as well. KDs with FDR < 15% were considered significant.

### Animal experiments

*LoxP*-floxed *Bckdk* (GenBank: NM_009739.3) exons 2-8 (*Bckdk ^f/f^*) strain and *LoxP*-floxed *Bckdha* (GenBank: NM_007533.5) exon 4 (*Bckdha ^f/f^*) strain (C57BL/6 background) were all generated by Cyagen Biosciences (Suzhou, China). Briefly, correctly targeted embryonic stem (ES) clones were confirmed via Southern Blotting and used for blastocyst microinjection, followed by chimera production. Founders were confirmed as germline-transmitted via crossbreeding with wild-type mice. F1 heterozygous mutant mice were confirmed as the final deliverables for this project. Fatty acid-binding protein 4–Cre recombinase (*Ap2-*cre) transgenic mice expressing Cre recombinase under the control of mouse *Fabp4* promoter (C57BL/6 background) were from Cyagen Biosciences (Suzhou, China). Adiponectin–Cre recombinase (*Adq-*cre) mice (C57BL/6 background) expressing Cre recombinase under the control of mouse *adiponectin* (*Adipoq*) promoter/enhancer regions were from The Jackson Laboratory. The *Bckdk ^f/f^* or *Bckdha ^f/f^* mice were crossed with the *Ap2-*cre strain to obtain *Ap2*-*Bckdk*-AKO or *Ap2*-*Bckdha*-AKO mice, in which the floxed *Bckdk* or *Bckdha* allele in adipose tissue was deleted, respectively. Similar strategy was applied with the *Adipoq*-Cre mice to obtain *Adq*-*Bckdk*-AKO or *Adq*-*Bckdha*-AKO mice. All studies were performed with male mice. All animals were housed at 22°C with a 12-hour light, 12-hour dark cycle with free access to water and standard chow.

To assess the development of obesity, high fat diet (Research diets, D12492, rodent diet with 60% kcal% fat) or normal chow diet (Research diets, D12450B, rodent diet with 10% kcal% fat) were used to feed mice at 6 weeks of age for different time. Body weight and food consumption were evaluated and body composition was measured by MRI (EchoMRI). Metabolic rates and physical activity were measured by a comprehensive lab animal monitoring system (Columbus Instruments). VO2, VCO2, and energy expenditure were measured. Mice were fasted for 6 hours before tissue harvest (8:00-14:00) to exclude metabolic changes due to immediate food intake. Tissue samples from white adipose tissue (epididymal fat and inguinal fat), brown adipose tissue (BAT), liver, and other tissues were weighted and quickly harvested and frozen in liquid nitrogen and maintained at −80°C until processed. All animal procedures were carried out in accordance with the guidelines and protocols approved by the Committee for Humane Treatment of Animals at Shanghai Jiao Tong University School of Medicine or the University of California at Los Angeles Institutional Animal Care and Use Committee (IACUC).

### Glucose and insulin tolerance test

After fasting for 6 hours (8:00-14:00), mice were intraperitoneal injected with D-glucose (1.5 g/kg body weight; Sigma, USA) for glucose tolerance test or injected with bovine insulin (0.75 U/kg bodyweight; Sigma, USA) for insulin tolerance test. Blood glucose was measured by OneTouch UltraEasy glucometer through tail bleeding at the times indicated after injection.

### Hematoxylin-eosin staining and adipocyte size counting

At the time of sacrifice, adipose tissues from mice were dissected and fixed with 4% paraformaldehyde. The fixed tissue was then dehydrated, cleared, and embedded in paraffin. Sections were stained with hematoxylin and eosin to assess morphology. For adipocyte size measurements, images were taken from each mouse (n=6-7 every group). AdipoCount software was used to measure cell size.

### Immunofluorescence

Immunostaining was performed to detect BrdU in paraffin sections according to the standard protocols. In brief, the fixed tissue was dehydrated, cleared, and embedded in paraffin. Sections were incubated with BrdU antibody (1:500 dilutions, GB12051, Servicebio) at 4℃ overnight, followed by PBS with corresponding secondary antibodies. For immunocytochemistry, the Hela cells were cultured on glass coverslips in 6-well plates for 24-48 hours before indicated treatments. After BCAA starvation and Leucine or DMAA treatment, cells on glass coverslips were washed once with PBS, fixed for 10 min with 4% paraformaldehyde, washed 3 times again with PBS, and permeabilized with 0.1% Triton X-100 in PBS. Cells were then washed 3 times, and blocked with 3% BSA in PBS for 30 min at room temperature. Cells on coverslips were then incubated in PBS with primary antibodies (1:200 dilutions, mTOR, CST #2983; 1:400 dilutions, LAMP2, SCBT sc-18822) at 4 °C overnight. Cells were then washed again, and incubated with Alexa Fluor 488 (A32723, Thermo) or Alexa Fluor 568 (A11011, Thermo) secondary antibodies in PBS for 1 hour at room temperature. After washing with PBS, coverslips were mounted to glass slides with a mounting buffer. Images were obtained on DeltaVision™ OMX SR super-resolution microscope. The colocalization was measured using Volocity software for Pearson’s correlation coefficient (PCC).

### Reagents

BCAA-free DMEM was customized from Invitrogen, USA. DMEM without methionine and cysteine was purchased from GIBCO (21012026). BCAAs, BCKAs, DMAA (3,3-Dimethylacrylic acid, D138606), (E)-3-Methylglutaconic acid (44108), HMGA (H4392), spermidine (S0266), spermine (S4264), N-(3-aminopropyl cyclohexylamine (APCHA, C108057), DMFO, methionine and cysteine were purchased from Sigma (USA). SAM486A was purchased from MedKoo (200411, USA). Rapamycin was obtained from ApexBio Technology (A8167).

### Isolation of primary white adipose mature adipocytes and SVF cells for differentiation

Subcutaneous (inguinal) white fat pads of 6 weeks age male mice were dissected, finely minced, and incubated with collagenase type IV (GIBCO17104, 10 mg/ml) in the isolation buffer (1% BSA, PBS [pH 7.4]) for 40 min at 37 °C on a shaker. At the indicated time, digested tissues were resuspended with twice volume of complete medium (DMEM supplemented with 10% FBS and 1% Penicillin/Streptomycin), followed by centrifuging at 800 g for 8 min to separate the SVF and the mature adipocytes fraction. Mature adipocytes were transferred to a new 5 ml centrifuge tube, and centrifuged at 800 g for 8 min, then sucked off the solution at the bottom of the centrifuge tube through narrow-bore micropipette tips and quickly frozen with liquid nitrogen and maintained at −80°C until processed. Pelleted SVF was resuspended in 2 ml DMEM supplemented with 10% FBS and 1% P/S and centrifuged at 1000 g for 8 min. Finally, the supernatant was discarded and pelleted SVF was quickly frozen with liquid nitrogen and maintained at −80°C until processed. For SVF cells differentiation, SVF cells were seeded on 6 cm dishes with 10% CS DMEM and incubated at 37℃ and 10% CO2. At 2 days post-confluence (day 0), cells were treated with the medium supplemented with 0.5 mM 3’-Isobutyl-1-methylxanthine (IBMX) (Sigma, Cat. I7018-1G), 1 µM dexamethasone (Sigma, Cat. D4902-25MG), 10 µg/ml insulin (Sigma, Cat. I5500-50MG), 0.2 nM T3 (Sigma) and 10 µM rosiglitazone (71740, Cayman). Two days later, cells were changed to medium containing insulin, T3 and rosiglitazone. The medium was replaced by fresh one every 2 days.

### Single cell RNA-seq

SVF preparation. *Bckdk^f/f^* and *Ap2***-***Bckdk***-**AKO (n=3 per group) at 6 weeks of age were fed 60% HFD for 2 weeks, and SVF cells were prepared as described above.

Single cell transcriptome capturation. Cells were firstly stained with two fluorescent dyes, Calcein AM (Thermo Fisher Scientific Cat. No. C1430) and Draq7 (Cat. No. 564904), for precisely determination of cell concentration and viability via BD Rhapsody™ Scanner. BD Rhapsody Express system was utilized for single cell transcriptomics capturation. Cells were loaded into one BD Rhapsody™ Cartridge which was primed and treated strictly following the manufacturers protocol (BD Biosciences). Cell Capture Beads (BD Biosciences) were then loaded excessively onto the cartridge to ensure that nearly every microwell contains one bead, and the excess beads were washed away from the cartridge. After lysing cells with lysis buffer (BD Biosciences), Cell Capture Beads were retrieved and washed prior to performing reverse transcription.

Library construction and sequencing. Briefly, double strand cDNA was firstly generated from microbeads-captured single cell transcriptome through several steps including reverse transcription, second strand synthesis, end preparation, adapter ligation and whole transcriptome amplification. Then final cDNA library was generated from double strand full length cDNA by random priming amplification with BD Rhapsody cDNA Kit (BD Biosciences, Cat. No. 633773) and BD Rhapsody Targeted mRNA & AbSeq Amplification Kit (BD Biosciences, Cat No. 633774). All the libraries were sequenced in a PE150 mode (Pair-End for 150bp read) on the NovaSeq platform (Illumina).

Data analysis. Raw sequencing reads of the cDNA library were processed through the BD Rhapsody Whole Transcriptome Assay Analysis Pipeline, which included filtering by reads quality, annotating reads, annotating molecules, determining putative cells and generating single-cell expression matrix. Among all the output files, Matrix for UMI counts per cell corrected by RSEC result were used for downstream clustering analysis.

The R package Seurat analysis. Raw gene expression matrices from the cartridge were read into R (version 3.6.3) and converted to Seurat objects. We filtered low quality cells that have unique feature counts less than 200, and that with more than 20% mitochondrial counts. Out of 17346 cells, 16969 cells remained. The gene expression matrix was then normalized to the total cellular UMI count. Top 2000 features were selected as highly variable genes for further clustering analysis. In order to reduce dimensionality, PCA was performed based on the highly variable genes after scaling the data with respect to UMI counts. On top of that, the first 50 principal components were chosen for downstream clustering based on heatmap of principle components, Jackstraw plot, and elbow plot of principle components to further reduce dimensionality using the tSNE algorithm. The transcriptional markers of each cluster were calculated using the FindAllMarkers function with the Wilcoxon Test under the following criteria: log2 fold change > 0.25; min. pct > 0.25. Top markers of each cluster were then selected to perform heatmap plot.

Unbiased single cell pseudotime analysis. After celltype annotation, we performed single cell pseudotime analysis for each celltype separately using Monocle2 with DDR-Tree reduction method (http://cole-trapnell-lab.github.io/monocle-release/). As for none-immune cell, single cell pseudotime analysis was performed with default parameters to eliminate batch effect. Briefly, raw gene expression matrix of none-immune cell was converted into a Monocle object. During feature selection, we chose ordering genes for downstream analysis only when their mean expression was larger than 0.1. Batch effect was also eliminated during dimensionality reduction. Trajectory plots, gene kinetics plots and heatmap were utilized to show pseudotime results.

Single Cell preparation, library construction, sequencing and analysis were performed by Shanghai Sinomics Corporation.

### Flow Cytometry

iWAT SVF cells were isolated from male mice treated with HFD for 2 weeks. The SVF cells were stained with LIVE/DEAD Fixable Viability Stain 780-APC-Cy^TM^7 (FVS780, 1:1000, 565388, BD Horizon) at 4℃ for 20min and then washed with stain buffer (554656, BD Pharmingen^TM^) containing FBS. SVF cells were stained with surface fluorescence antibodies including CD45-BV510 (1:200, 563891, BD Pharmingen), CD36-BB515 (1:200,565933, BD Pharmingen), PDGFRA (CD140a)-BB700 (1:200, 742176, BD Pharmingen) at 4℃ for 40min, and washed with stain buffer. For intracellular antibody staining, SVF cells were fixed/permeated with fixation/permeabilization kit (554714, BD Pharmingen^TM^) at 4℃ for 30min and then washed with stain buffer. Subsequently, cells were stained with PPARr (1:200, NBP2-76958, Novus) and F(ab’)2-donkey anti-Rabbit IgG (H+L) secondary antibody PE (1:100, 12-4739-81, BD bioscience) at 4℃ for 1h, washed with stain buffer. Finally, the SVF cells were incubated with Stain Buffer. All cytometric data were collected on FACS BD LSRFortessa X-20 cell analyzer. Cell population (%) was calculated as frequency for parent. Flow Jo software (version 10.6.2) was used for data analysis.

### Cell culture and 3T3-L1 cell differentiation

HEK 293FT cells, 3T3-L1 cells, and Hela cells were purchased from American Type Culture Collection (ATCC; Manassas, VA). Cells were maintained in Dulbecco’s Modified Eagle’s Medium (DMEM) containing 100 U/ml penicillin and 100 μg/ml streptomycin and 10% fetal bovine serum (FBS), and incubated in a humidified incubator at 37°C and 5% or 10% CO2. 3T3-L1 pre-adipocytes were cultured in DMEM supplemented with 4 mM l-glutamine, 4.5 g/l glucose, 8 μg/ml biotin, 0.11 g/l sodium pyruvate and 10% newborn calf serum (NCS), 100 U/ml penicillin and 100 μg/ml streptomycin, and incubated in a humidified incubator at 37°C and 10% CO2. After the cell confluence was 100%, a standard differential protocol was performed. 3T3-L1 cells were differentiated in a similar way as described above in SVF cells differentiation, but T3 and rosiglitazone was not included in the adipogenic cocktail. Differentiated 3T3-L1 adipocytes (Day 8) were washed with PBS and fixed in 4% paraformaldehyde for 5min. Oil Red O dye (Sigma) dissolved in isopropanol was used to stain the cells for 1 h. The images were recorded using an Olympus microscope or a digital camera. All experiments were repeated at least twice independently with similar results.

### shRNA lentivirus production and infection

shRNA-encoding plasmids were obtained from the Thermo Scientific Open Biosystems GIPZ Lentiviral shRNAmir Library. The hairpin vectors were co-transfected with the lentivirus expression plasmid and packaging plasmid into actively growing HEK293FT using X-tremeGENE 9 DNA transfection reagent (Roche) according to the manufacturer’s instructions. Virus containing supernatants were collected at 48 hours after transfection and filtered through 0.45 µm cellulose acetate filters. 3T3-L1 cells were plated in 60mm dishes at 20-30% confluence and infected for 24 hours in the presence of 4 µg/ml polybrene. After infection, the cells were split into fresh media for differentiation. Knockdown efficiency was confirmed by qPCR or western blot. BCAT2 shRNA: GTCACTATGAAGGAATTGA; IVD shRNA: AACAAGTTCTGGATCACCA; MCCC1 shRNA: AACTCCACACTCAAGATCA; AUH shRNA: AGCCATGTGTTAGAACAGA.

### Metabolomics analysis

During differentiating, on the day 4, 1 ml cell media were collected, flash-frozen and stored at −80℃. Then 3T3-L1 adipocytes were gentle trypsinized, and transferred to 15 ml polypropylene tubes, then spined at about 1000 g for 3 min to pellet cells. After centrifuging, supernatant was removed. Cell pellet were centrifuged at 1000 g for 3 min again to remove all supernatant, and then flash-frozen and stored at −80°C. The metabolomic analysis of 3T3-L1 cells and media were carried out by Metabolon, Inc. (Durham, NC). Samples were prepared using the automated MicroLab STAR® system from Hamilton Company. Several types of controls were analyzed in concert with the experimental samples. The LC-MS portion of the platform was based on a Waters ACQUITY ultra-performance liquid chromatography (UPLC) and a Thermo-Finnigan LTQ mass spectrometer operated at nominal mass resolution, which consisted of an electrospray ionization (ESI) source and linear ion-trap (LIT) mass analyzer. The samples destined for analysis by GC-MS were dried under vacuum prior to being derivatized under dried nitrogen using bistrimethyl-silyltrifluoroacetamide. Derivatized samples were separated on a 5% diphenyl / 95% dimethyl polysiloxane fused silica column and analyzed on a Thermo-Finnigan Trace DSQ fast-scanning single-quadrupole mass spectrometer using electron impact ionization (EI) and operated at unit mass resolving power. Raw data was extracted, peak-identified and QC processed using Metabolon’s hardware and software. Peaks were quantified using area-under-the-curve. A collection of information interpretation and visualization tools for use by data analysts. Welch’s two-sample t-test is used to test whether two unknown means are different from two independent populations.

### 3-methylcrotonyl-CoA measurement

Levels of 3-methylcrotonyl-CoA in 3T3-L1 adipocytes was analyzed by UHPLC-MS/MS. Briefly, extraction and homogenization of metabolites were done through cold methanol, 50 mM NH4HCO3 and 80% methanol (including 10 mM NH4HCO3). Then samples were processed by 5 cycles of 1 min ultra-sonication and 1 min interval in ice-water bath. After centrifugation at 4°C and 16000 g for 15 min, 200 μl supernatants were evaporated to dryness under nitrogen stream. The dry residues were reconstituted in 50 μl of 50% acetonitrile containing 5 mM ammonium acetate and 1 μg /ml malonyl-^13^C3-CoA. After centrifugation at 16000 g and 4°C for 15 min, the supernatant was utilized to perform UHPLC-MS/MS analysis. The quality control (QC) sample was obtained by pooling all the prepared samples. The UHPLC-MS/MS analysis was performed on an Agilent 1290 Infinity II UHPLC system coupled to a 6470A Triple Quadrupole mass spectrometry (Santa Clara, CA, United States). The eluted analysts were ionized in an electro spray ionization source in positive mode (ESI+). The raw data were processed by Agilent MassHunter Workstation Software (version B.08.00) by using the default parameters and assisting manual inspection to ensure the qualitative and quantitative accuracies of each compound. The peak areas of target compounds were integrated and output for quantitative calculation. The analysis was carried out by Shanghai Profleader Biotech Ltd, China.

### Measurement of polyamine synthesis from methionine

Sample Preparation: 3T3-L1 adipocytes on the 4th day of differentiation were incubated with L-methionine-^13^C5,^15^N (30ug/mL) in a methionine-deficient DMEM for 3.5 hours. Cells were then washed with PBS and collected in 80% aqueous methanol were processed by ultrasonication in ice-water for 3 min. The cell lysates were centrifuged at 15000 g and 4℃ for 15 minutes, and the supernatant was dried under nitrogen stream. The residue was reconstituted in 20 μL of 50% acetonitrile containing 10 μg/ml L-arginine-^13^C6-^15^N4 as internal standard. The chemical derivatization was performed at room temperature for 10 minutes after adding 40 μL of sodium carbonate (100mM) and 40 μL of 2% benzoyl chloride in acetonitrile. The derivatives were centrifuged and applied to analysis of UHPLC-HRMS. Metabolite Measurement by UHPLC-HRMS: Chromatographic separation was performed on a ThermoFisher Ultimate 3000 UHPLC system with a Waters BEH C18 column (2.1mm × 100 mm, 1.7 μm). The injection volume was 3μL and the flow rate was 0.25 mL/min. The column temperature was 50°C. The mobile phases consisted of water with 0.1% formic acid (phase A) and methanol with 0.1% formic acid (phase B). A linear gradient elution was performed with the following program: 0 min,2%B; 9 min, 70% B;14min, 100%B and held to 18 min; 18.1 min, 2%B and held to 20 min. The eluents were analyzed on a ThermoFisher Q Exactive™ Hybrid Quadrupole-Orbitrap™ Mass Spectrometry (QE) in Heated Electrospray Ionization Positive (HESI+) mode, separately. Spray voltage was 4000 V. Both Capillary and Probe Heater Temperature were set to 320 °C. Sheath gas flow rate was 35 (Arb, arbitrary unit), and Aux gas flow rate was 10 (Arb). S-Lens RF Level was 50 (Arb). The full scan was operated at a high-resolution of 70000 FWHM (m/z=200) at a range of 70- 1050 m/z with AGC Target setting at 3×106.

Data Analysis: Raw data was preprocessed by Xcalibur (v4.0), and the peak areas were applied to correct natural isotope and calculate mass isotope distribution (MID) by IsoCor (Bioinformatics, 2019, doi: 10.1093/bioinformatics/btz209).

### Western blot analysis

Whole cell lysate or tissues were extracted using lysis buffer (250 mM Tris-HCl [pH6.8], 2%SDS, glycerol 10% and protease/phosphatase inhibitors cocktail). Protein concentration was determined using BCA Protein Assay Kit (TIANGEN) and diluted in loading dye (β-mercaptoethanol 5%, 0.02% bromophenol blue), heated at 99°C for 5 min, and separated by SDS-PAGE, and transferred onto NC membranes. Membranes were blocked with 2% non-fat milk (BD) in Tris-buffered saline (TBS) containing 0.1% Tween 20 (TBST), probed with the indicated primary antibodies at 4°C overnight. Protein signals were detected using HRP conjugated secondary antibodies and enhanced chemiluminescence (ECL) western blotting detection regents (Pierce). The BCKDK antibody (AV52131) and AUH antibody (AV40374) were purchased from Sigma. Anti-PPARγ (#81B8), anti-AKT (#9272), anti-AKT^S473^ (#4060), anti-AKT^T308^ (#2965), anti-p70S6K (#9202), anti-p70S6K^T389^ (#9234), anti-Erk1/2 (#9102) antibodies were purchased from Cell Signaling Technology. Anti-C/EBPα (ab18336), anti-C/EBPβ (ab40764) and anti-MCCC1 (ab174984) antibodies were purchased from Abcam. Anti-BCKDE1α (sc-271538), anti-BCKDE1β (sc-374630), anti-BCAT2 (sc-169986) and anti-IVD (sc-514240) antibodies were purchased from Santa Cruz Biotechnology. Anti-β-actin (AB2001) and anti-ODC1 (AY3310) were purchased from Abways Technology. Anti-SAT1 antibody (NB110-41622) was from Novus Biologicals and anti-AMD1 antibody (11052-1-AP) was from Proteintech.

### RNA isolation and qPCR

Total RNA was extracted using the Trizol (Invitrogen, USA) according to the manufacturer’s instructions. Total RNA (1.5 μg) was reverse transcribed using random primers and MMLV (Promega, USA). Each cDNA sample was analyzed in triplicate with the Applied Biosystems Prism7900HT Real-Time PCR System using Absolute SYBR Green (ABI, USA). The primer sequences were listed in (Supplementary Table 3).

### Measurement of amino acid content by LC-MS/MS

Calibration standards were prepared at concentrations of 5, 10, 25, 50, 100, 250 and 500 ng/ml by diluting a fixed amount of stock solution of glutamate in KHB. The solution with known spiked amount of amino acid was defined as the quality control (QC), which was set at low, middle, and high concentrations (10, 100, and 400 ng/ml). The QC samples were prepared independently from the calibration samples. The stable isotope-labeled internal standard amino acids were used as the IS in the calibration standards and QC samples, and the concentration was kept consistent at 40 ng/ml. All calibration standards and QC samples were stored at −80°C until the LC-MS/MS analysis.

One hundred microliters of sample were mixed with 100 μl of methanol, containing IS at a concentration of 40 ng/ml. The mixture was vortex mixed for 30 sec and centrifuged at 13000 g for 10 min. Two microliters of the supernatant were analyzed using LC-MS/MS.

Liquid chromatography was performed using an Agilent 1200 HPLC system (Agilent Technologies, CA, USA), and separation was carried out at 40°C using a ZIC-HILIC column (2.1 mm × 100 mm, 3 μm; Merck Sequant, Umea, Sweden). Isocratic elution was performed. An ESI in positive ionization mode was used. The mode of multiple reaction monitoring (MRM) was used to identify and quantify amino acids. The analytic data were processed using the MassHunter software package (Agilent Technologies, CA, USA), which contained qualitative and quantitative analysis modules.

### Statistics

Unless otherwise specified, statistical analyses were performed with two-sided Student’s t-test (two groups) and one-way (> two groups) or two-way (tolerance tests) ANOVA, followed by a Tukey post hoc test where appropriate using GraphPad Prism. Data were calculated as the mean±SEM (standard error of the mean) unless otherwise indicated. A *p* value of less than 0.05 was considered statistically significant.

## ACKNOWLEDGEMENTS

This work was supported by the National Key Research and Development Program of China (2019YFA0802503), the National Natural Science Foundation of China (92057107, 81570717, 31900819), Collaborative Innovation Program of Shanghai Municipal Health Commission (2020CXJQ01), Tianjin Key Medical Discipline (Specialty) Construction Project (TJYXZDXK-032A), and the Science and Technology Commission of Shanghai Municipality (16JC1404400).

## AUTHOR CONTRIBUTIONS

H. Sun designed the project and directed the research; J. Shao, Y. Liu, X. Zhang, L. Shu, J. Yu, S. Yang, C. Wang, N. Cao, M. Zhou, R. Chi, J. Wang, M. Chen, C. Liu, and J. Wang performed the research; J. Shao, Y. Liu, X. Zhang, J. Yu, L. Shu, C. Gao, M. Zhou, J. Wang and H. Sun analyzed the data; R. Liu, J. Wang, Y. Wang, W. Zhang, G. Ning, W. Wang, and X. Yang helped to design the overall study and analyzed the data; all authors contributed to the manuscript preparation.

## Supplementary Materials

**Fig. S1.**
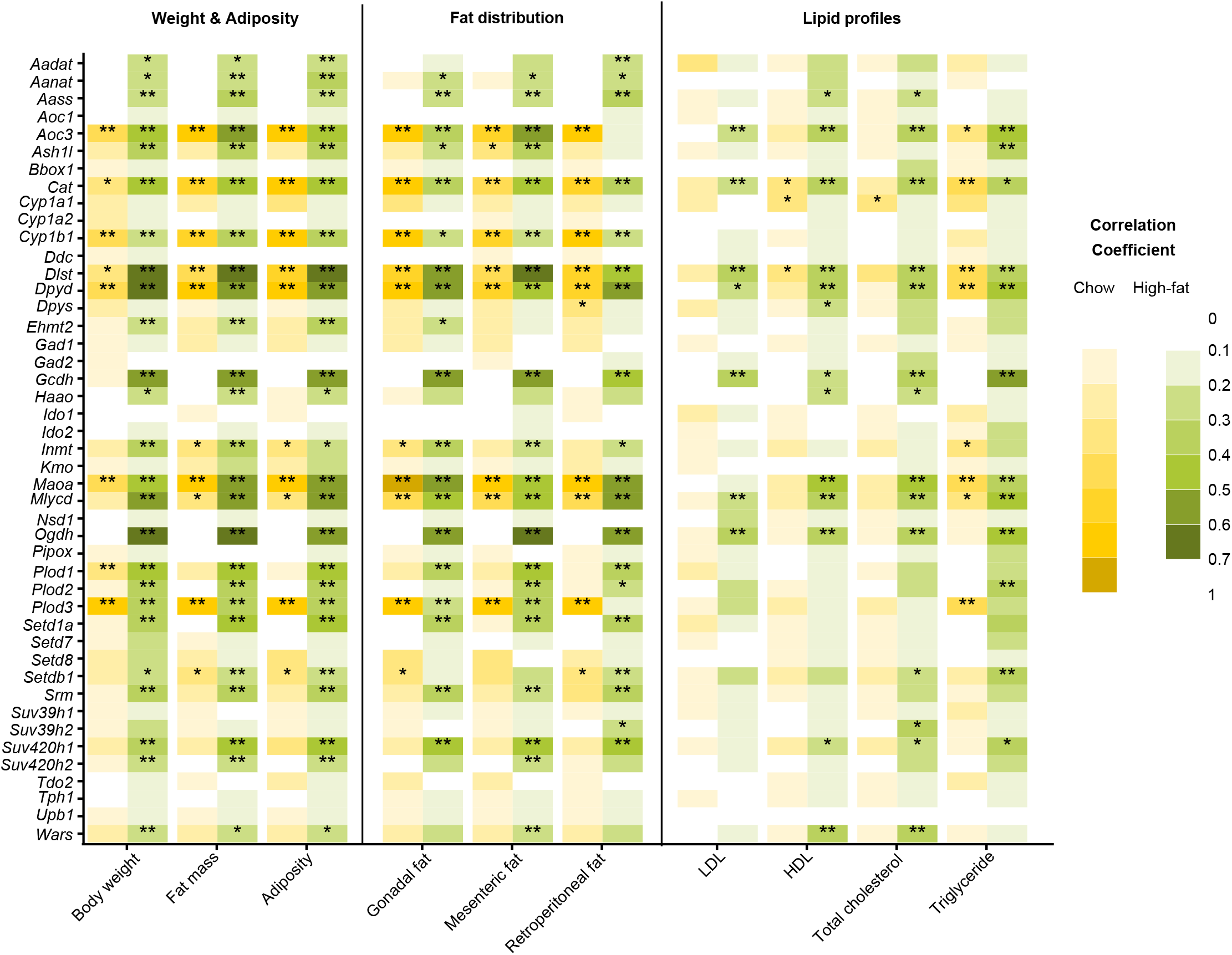
Gene-trait correlation for genes in non-BCAA amino acids metabolic pathways in the adipose tissue from HMDP mice fed with chow diet and high-fat diet. *passing Bonferroni p < 0.05 corrected for the number of BCAA genes; **passing Bonferroni p < 0.05 corrected for the numbers of BCAA genes and traits.

**Fig. S2.**
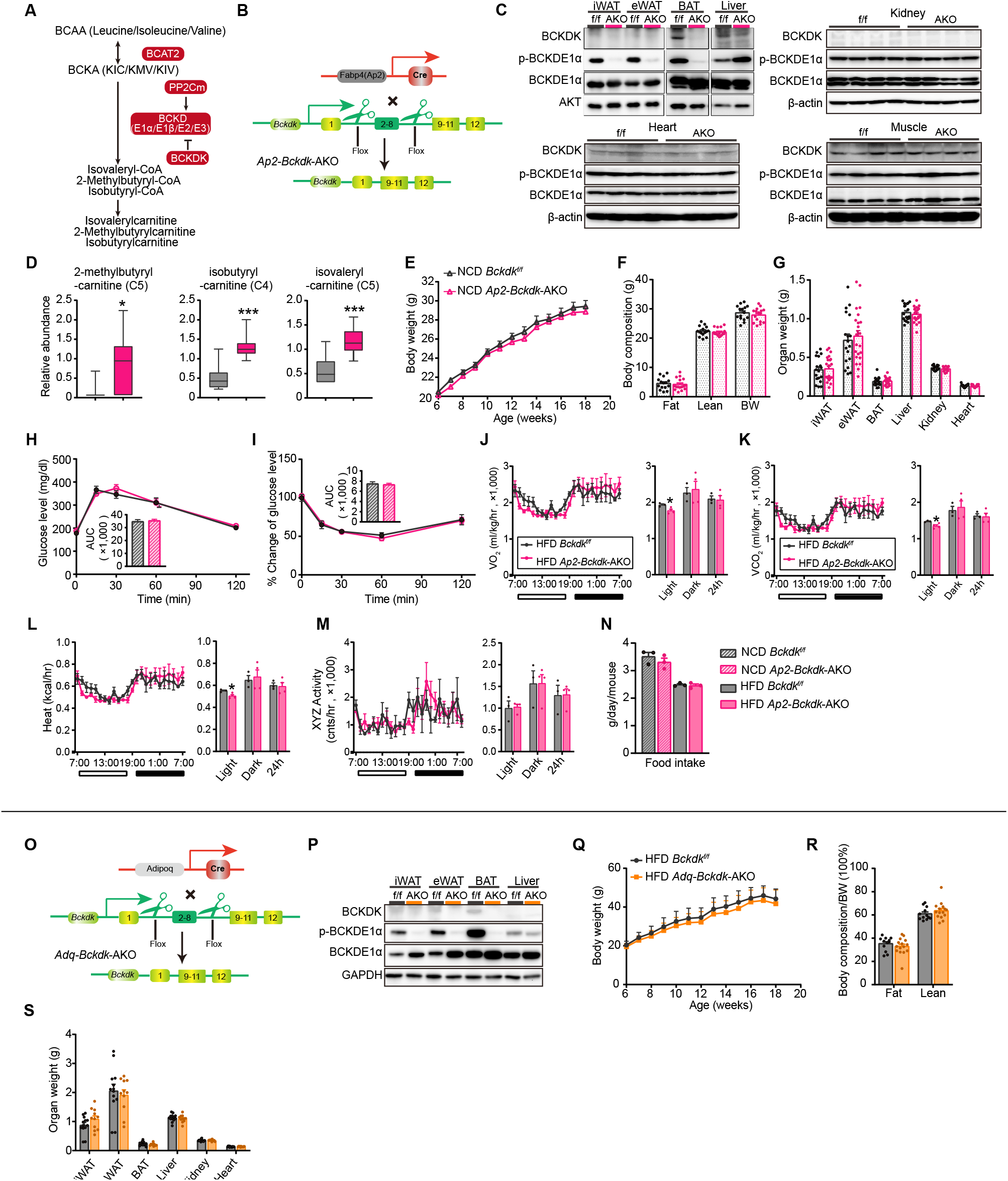
Characterization of *Ap2*-*Bckdk*-AKO and *Adq-Bckdk*-AKO mice. (A) Illustration of partial BCAA catabolic pathway with enzymes, intermediates, and derivatives. (B) Schematic of conditional *Bckdk* deletion in adipose tissue with *Fabp4* (*Ap2*)-Cre. (C) Western blot analysis of BCKDK, p-BCKDE1α, BCKDE1α in the indicated tissues from f/f (*Bckdk^f/f^*) and AKO (*Ap2-Bckdk*-AKO) mice. (D) Metabolomic analysis of BCAA catabolites in the sWAT of *Bckdk^f/f^* and *Ap2*-*Bckdk*-AKO mice. (E) Body weight (n=15- 18 per group) of male *Bckdk^f/f^* and *Ap2-Bckdk*- AKO mice on normal chow diet (NCD) for 12 weeks. (F-I) Body composition (n=9-12 per group) (F), organ weight (n=15-16 per group) (G), glucose tolerance test (GTT) (n=15 per group) and the area under the curve (AUC) (H), insulin tolerance test (ITT) (n=8-12 per group) and the area under the curve (AUC) (I) of male *Bckdk^f/f^* and *Ap2-Bckdk*-AKO mice on normal chow diet (NCD) for 6 weeks. (J-M) Oxygen consumption (VO2) (J), carbon dioxide production (VCO2) (K), heat (M), and locomotor activity (L) from 2 weeks HFD-fed male *Bckdk^f/f^* and *Ap2-Bckdk*-AKO mice (n=3- 4 per group). (N) Food intake in *Bckdk^f/f^* and *Ap2-Bckdk*-AKO mice fed NCD (n=12-15 per group) and HFD (n=13 per group). Schematic of conditional *Bckdk* deletion in adipose tissue with *Adipoq*-Cre. (P) Western blot analysis of BCKDK, p-BCKDE1α, BCKDE1α in the indicated tissues from f/f (*Bckdk^f/f^*) and AKO (*Adq-Bckdk*-AKO) mice. (Q-S) Body weight (n=12-15 per group) (Q), body composition (n=12-15 per group) (R), and organ weight (n=11-13 per group) (S) of male *Bckdk^f/f^* and *Adq-Bckdk*-AKO mice on high fat diet (HFD) for 12 weeks. Data are represented as mean ± s.e.m. Statistical significance was determined by unpaired two-tailed Student’s t-test or two-way analysis of variance (ANOVA). *p<0.05, ***p<0.001.

**Fig. S3.**
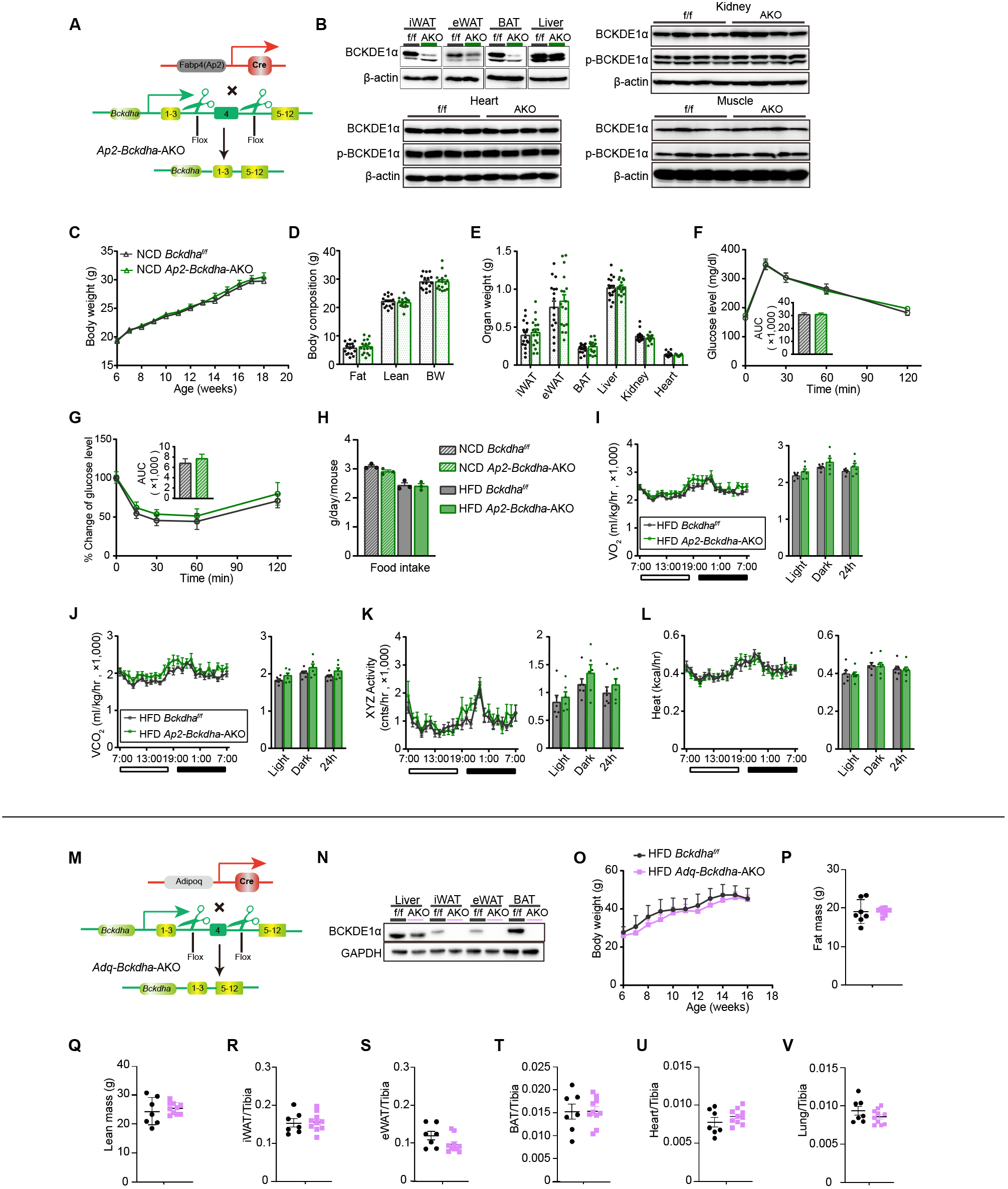
Characterization of *Ap2*-*Bckdha*-AKO and *Adq-Bckdha*-AKO mice. (A) Schematic of conditional *Bckdha* deletion in adipose tissue with *Fabp4* (*Ap2*)-Cre. (B) Western blot analysis of BCKDE1α in the indicated tissues from f/f (*Bckdha^f/f^*) and AKO (*Ap2-Bckdha*-AKO) mice. (C) Body weight (n=15-19 per group) of male *Bckdha^f/f^* and *Ap2-Bckdha*-AKO mice on normal chow diet (NCD) for 12 weeks. (D-E) Body composition (n=16-18 per group) (D), and organ weight (n=13-14 per group) (E) of male *Bckdha^f/f^* and *Ap2-Bckdha*-AKO mice on normal chow diet (NCD) for 6 weeks. (F-G) Glucose tolerance test (GTT) (n=15 per group) and the area under the curve (AUC) (F), insulin tolerance test (ITT) (n=11-18 per group) and the area under the curve (AUC) (G) of male *Bckdha^f/f^* and *Ap2-Bckdha*-AKO mice on normal chow diet (NCD) for 2 weeks. (H) Food intake in *Bckdha ^f/f^* and *Ap2-Bckdha*-AKO mice fed NCD (n=12 per group) and HFD (n=12 per group). (I-L) Oxygen consumption (VO2) (I), carbon dioxide production (VCO2) (J), locomotor activity (K), and heat (L) from 4 weeks HFD-fed male *Bckdha^f/f^* and *Ap2-Bckdha*-AKO mice (n=6 per group). (M) Schematic of conditional *Bckdha* deletion in adipose tissue with *Adipoq*-Cre. (N) Western blot analysis of BCKDE1α in the indicated tissues from f/f (*Bckdha^f/f^*) and AKO (*Adq-Bckdha*-AKO) mice. (O) Body weight (n=12-13 per group) of male *Bckdha^f/f^* and *Adq-Bckdha*-AKO mice on high fat diet (HFD) for 10 weeks. (P-Q) Body composition: fat mass (P), lean mass (Q) (n=7-9 per group male) of *Bckdha^f/f^* and *Adq-Bckdha*-AKO mice on HFD for 10 weeks. (R-V) Organ weight: iWAT (R), eWAT (S), BAT (T), heart (U), and lung (V) (n=7-10 per group male) of *Bckdha^f/f^* and *Adq-Bckdha*-AKO mice on HFD for 12 weeks. Data are represented as mean ± s.e.m. Statistical significance was determined by unpaired two-tailed Student’s t-test or two-way analysis of variance (ANOVA).

**Fig. S4.**
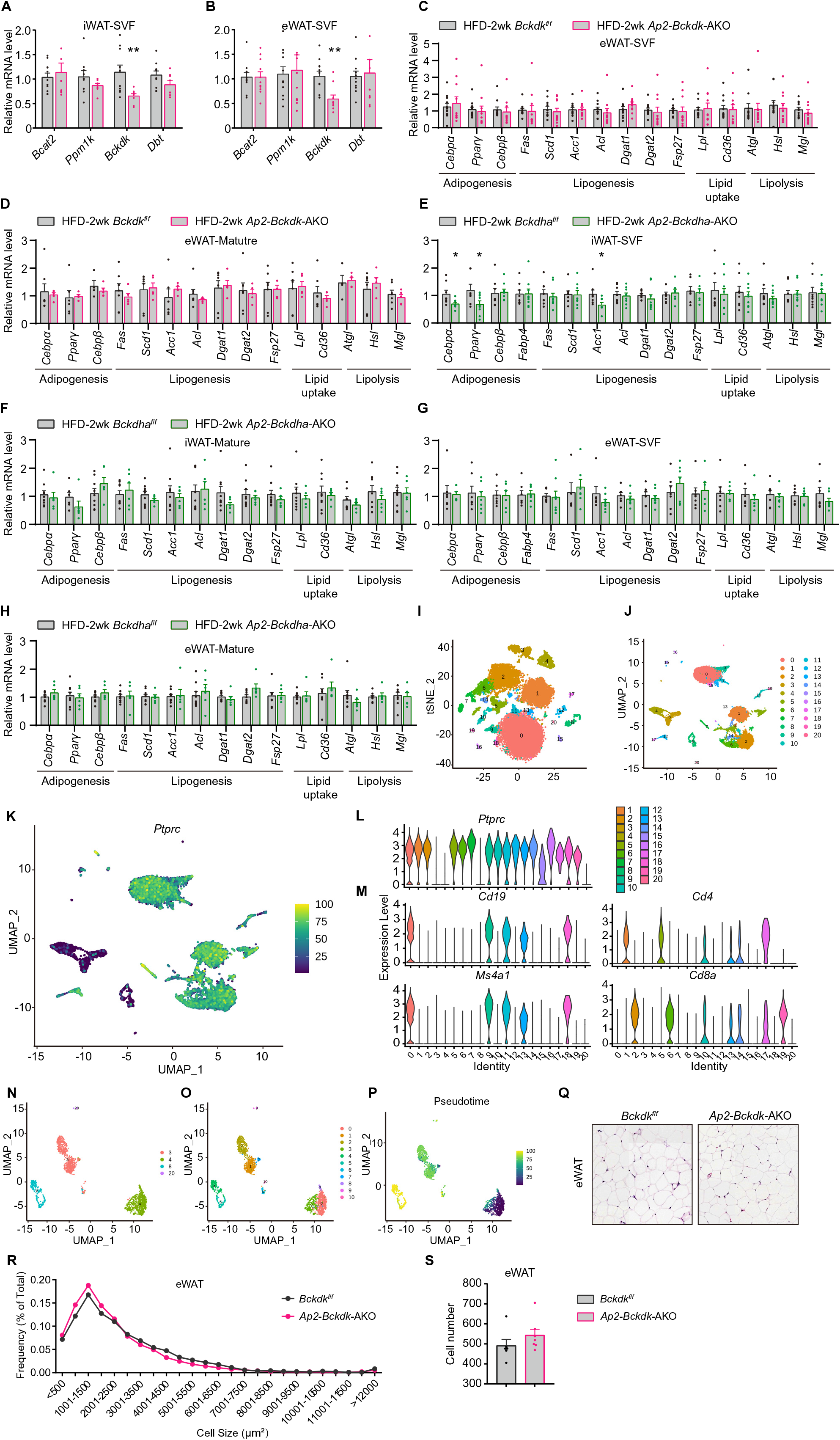
Gene expression in SVF and single cell sequencing analysis of SVF cells. (A-B) mRNA levels of BCAA catabolic genes in the SVF from the iWAT (A) and eWAT (B) of *Bckdk^f/f^* and *Ap2*-*Bckdk*-AKO mice (n=11 per group). (C-D) qPCR analysis of the expression of indicated genes in the SVF (n=11 per group) (C) and mature adipocytes (Mature) (n=4-5 per group) (D) of eWAT from male *Bckdk^f/f^* and *Ap2-Bckdk*-AKO mice on HFD for 2 weeks. (E-H) qPCR analysis of the expression of indicated genes in the SVF (n=7-9 per group) (E) and mature adipocytes (Mature) (n=6-9 per group) (F) of iWAT or SVF (n=5-9 per group) (G) and Mature (n=6-9 per group) (H) of eWAT from male *Bckdha^f/f^*and *Ap2-Bckdha*-AKO mice on HFD for 2 weeks. (I-S) Single cell sequencing analysis of SVF cells from iWAT of male Bckdk*^f/f^* and *Ap2*-*Bckdk*-AKO mice on HFD for 2 weeks (n=3 per group). T-distributed stochastic neighbor embedding (tSNE) plot (I) and UMAPs plot (J) of SVF cells; expression of leukocyte marker *Ptprc* in different clusters of SVF cells shown in the integrated UMAP plot (K) and violin plots (L); violin plots of canonical markers *Cd19*, *Cd4*, *Ms4a1*, *Cd8a* (M); 3, 4, 8 and 20 seurat clusters shown in the integrated UMAP plot (N); group 1-10 of CD45-SVF cells shown in the UMAP plot (O); feature plots of pseudotime analysis (P); histology(Q) and adipocyte size (R) and number (S) analysis in the eWAT of *Bckdk^f/f^* and *Ap2-Bckdk*-AKO mice on HFD for 2 weeks. Data are represented as mean ± s.e.m. Statistical significance was determined by unpaired two -tailed Student’s t-test. *p<0.05, **p<0.01.

**Fig. S5.**
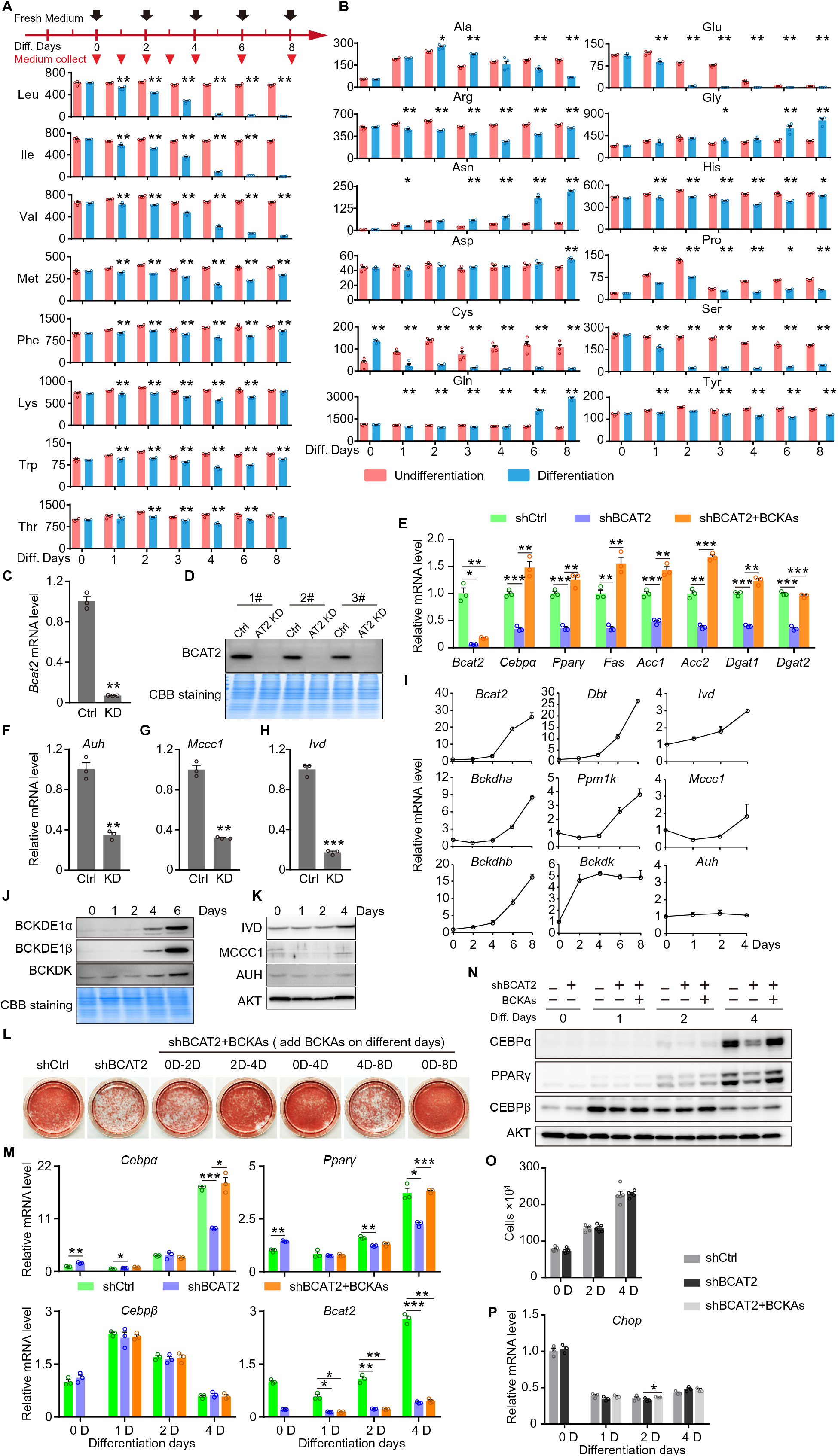
BCAA catabolism is essential for the differentiation of adipocytes. (A-B) Differentiation schema of 3T3-L1 cells (top) and individual essential amino acid concentrations (μM, y axis) in culture medium (A) and nonessential amino acid concentrations in culture medium (B) collected at different days during 3T3-L1 differentiation. The medium was replaced by fresh one every 2 days. * p<0.05, ** p<0.01 versus undifferentiated cells. (C-D) *Bcat2* mRNA (C) and protein (D) level in 3T3-L1 differentiated adipocyte with shRNA silencing. (E) qPCR results of specific gene in differentiated 3T3-L1 adipocytes with *Bcat2* silencing and BCKAs supplementation. (F-H) mRNA level of *Auh* (F), *Mccc1* (G), and *Ivd* (H) in 3T3-L1 differentiated adipocytes using shRNA silencing (KD). (I-K) qPCR (I) and western blot (J, K) results of BCAA catabolic genes using mRNA or lysates from 3T3-L1 cells at designated time points (0,1,2,4,6,8 days) of adipocyte differentiation. Coomassie brilliant blue (CBB) staining showed equal loading (H). (L) Images of oil red staining of differentiated 3T3-L1 adipocytes with shRNA silencing of BCAT2, treated with BCKAs during different periods of differentiation. (M-N) Specific gene mRNA level (M) and protein expression (N) in 3T3-L1 cells at designated time points of adipocyte differentiation with shRNA silencing of *Bcat2* and/or BCKAs supplementation. (O) Cells were counted at the indicated times of 3T3-L1 differentiation. (P) *Chop* mRNA levels at the indicated time. Data are represented as mean ± s.e.m. Statistical significance was determined by unpaired two-tailed Student’s t-test or one-way analysis of variance (ANOVA). *p<0.05, **p<0.01, ***p<0.001.

**Fig. S6.**
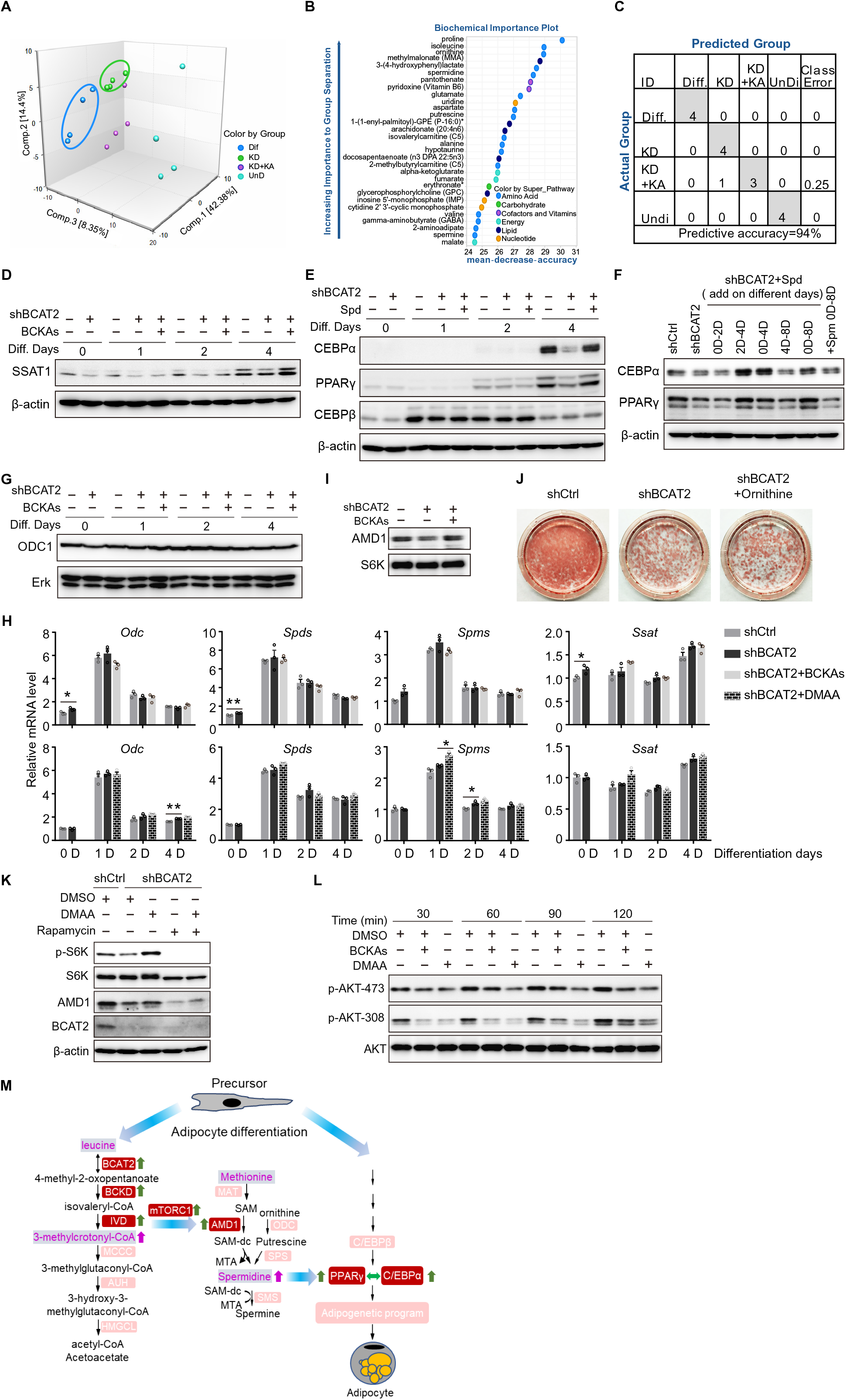
The leucine catabolite promotes differentiation through increasing mTORC1-AMD1-mediated polyamine synthesis. (A) Principal component analysis (PCA) of the metabolomics profiles visualized the experimental groups exhibiting different overall metabolic signatures. (B) Random Forest Confusion Matrix. (C) List of the top 30 biochemicals that separated different groups based on their importance, suggesting key differences in the metabolism of amino acids, cofactors and vitamins, lipids, and energy metabolism. (D) SSAT1 expression on different days in differentiating 3T3-L1 adipocytes with *Bcat2* silencing and/or BCKAs supplementation. (E-F) Protein expression of CEBPα, PPARγ and CEBPβ in differentiating 3T3-L1 adipocyte on different days with *Bcat2* silencing and spermidine (Spd) treatment. (G) ODC1 expression in differentiating 3T3-L1 adipocytes on different days. (H) Expression of polyamine metabolism related genes in differentiating 3T3-L1 adipocytes treated with BCKAs or DMAA and *Bcat2* shRNA. (I) AMD1 expression on the 4^th^ day of 3T3-L1 adipocyte differentiation. (J) Images of oil red staining of differentiated 3T3-L1 adipocytes with *Bcat2* silencing in the presence or absence of ornithine (5 mM). (K) Western blot analysis of AMD1 and mTOR signaling in differentiating 3T3-L1 adipocytes with *Bcat2* silencing, treated with DMAA with or without rapamycin for 6 hours. (L) Western blot analysis of AKT pathway in undifferentiated 3T3-L1 cells with BCKAs or DMAA treatment in the absence of BCAAs in culture medium. (M) Working model. BCAA catabolism promotes adipogenesis via leucine metabolite. Data are represented as mean ± s.e.m. Statistical significance was determined by unpaired two-tailed Student’s t-test or one-way analysis of variance (ANOVA). *p<0.05, **p<0.01.

**Table S1.**
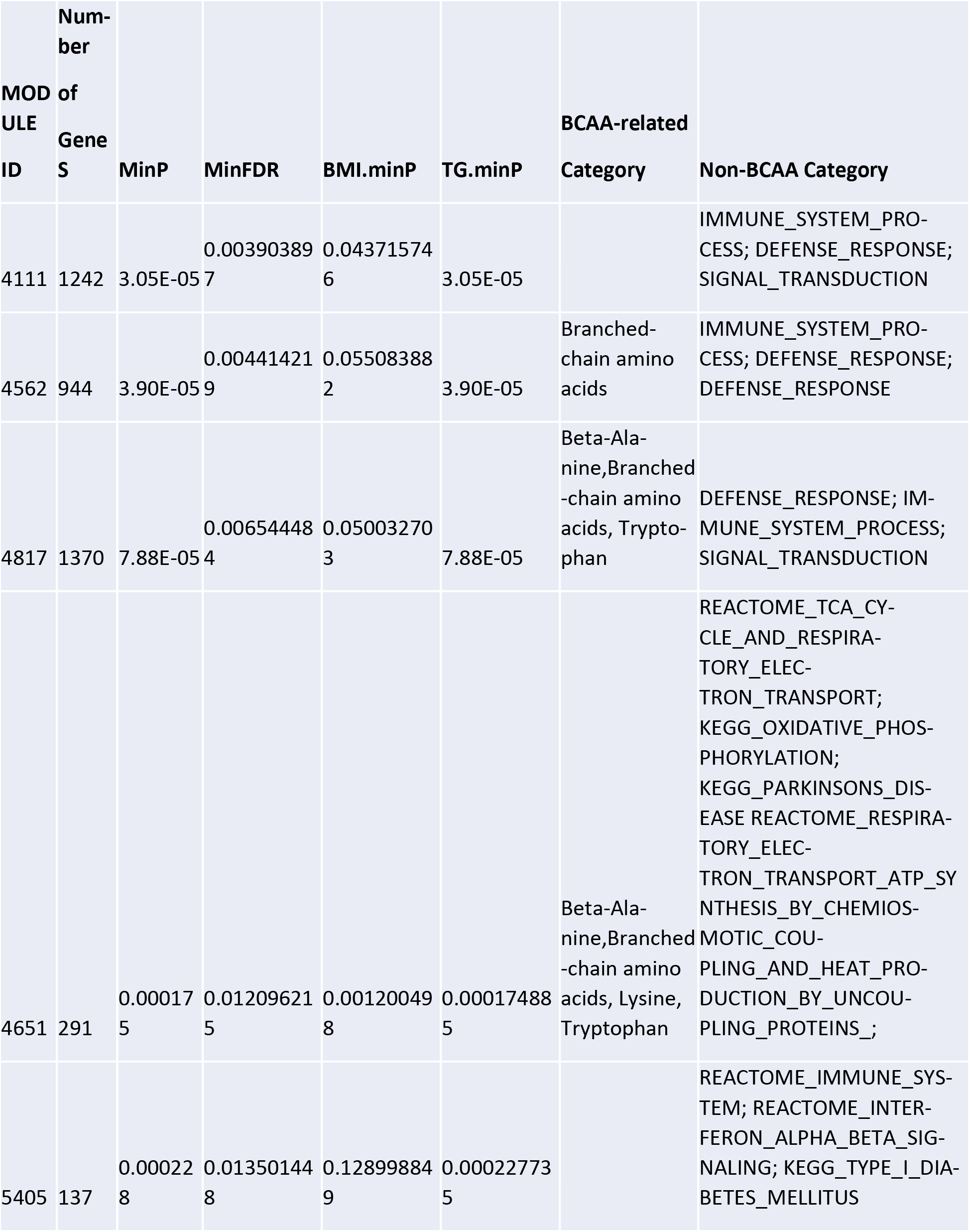
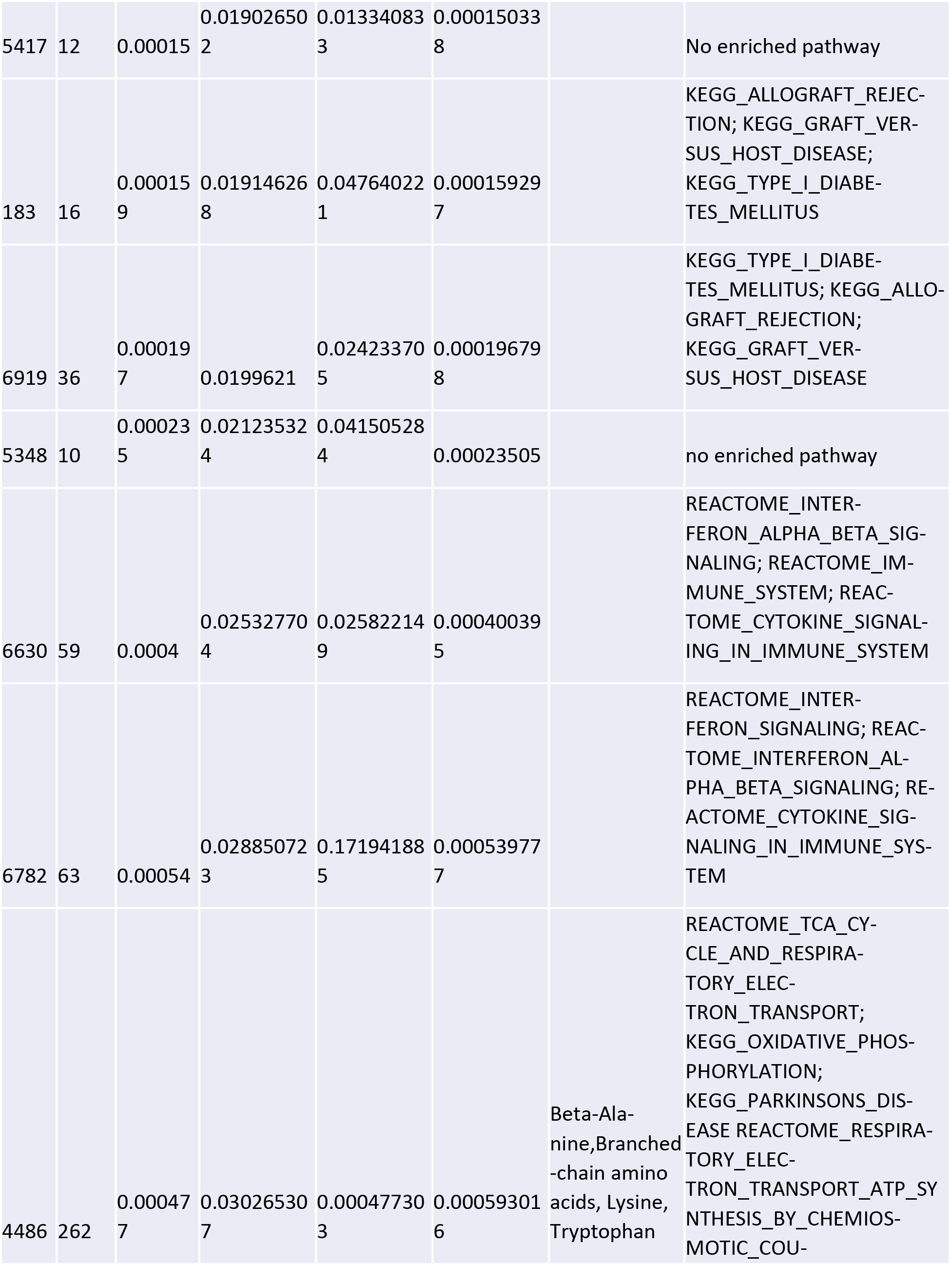
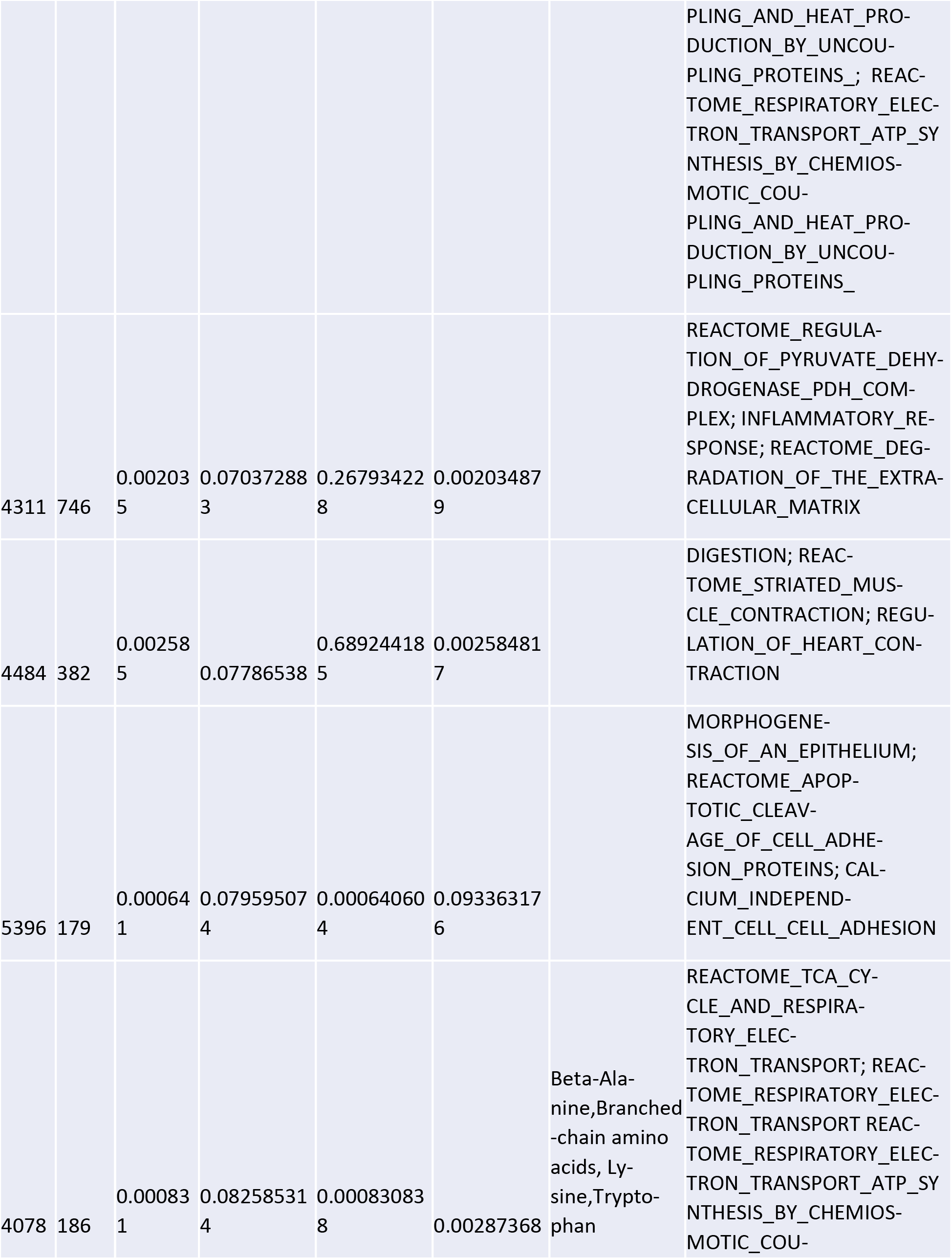
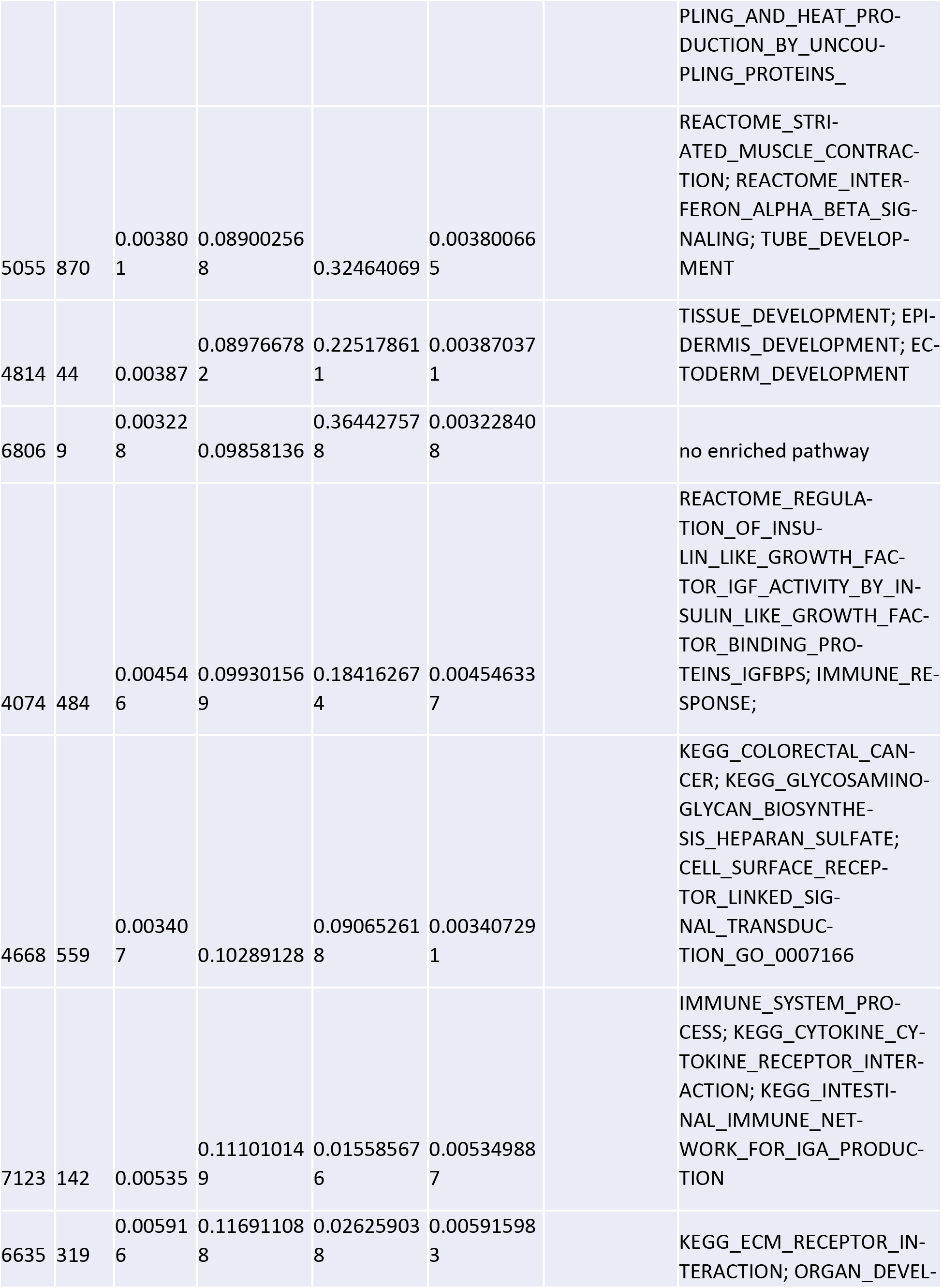
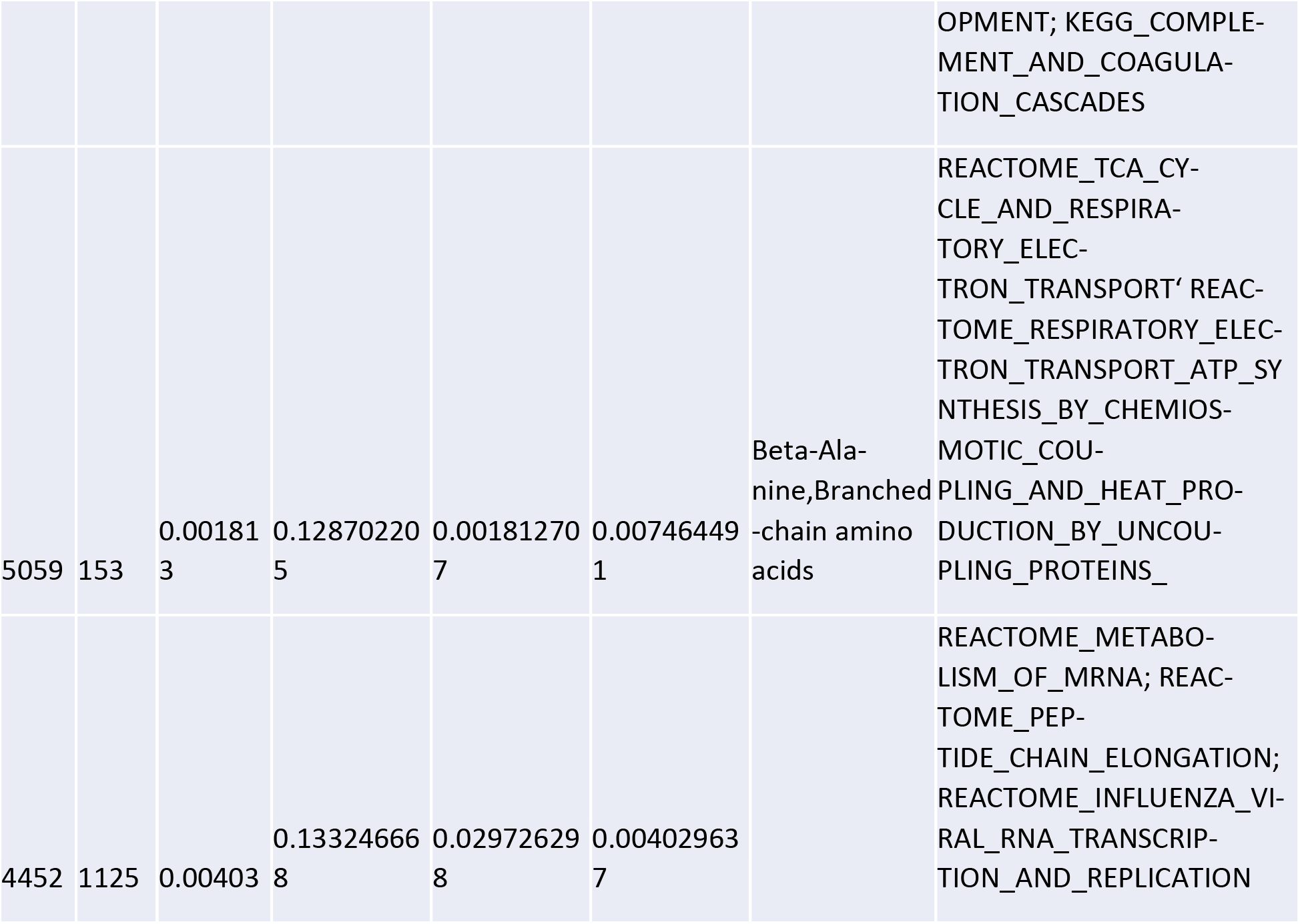
Enrichment of BCAA genes and non-BCAA KEGG pathways in the top of 25 co-expression modules associated with human clinical traits.

**Table S2.** Sources of adipose coexpression modules, eQTLs and Bayesian networks.

| Species | eQTL | Coexpression modules | Bayesian networks | Dataset |
| --- | --- | --- | --- | --- |
| Human | yes | yes | yes | 1,675 individuals from two Icelandic cohorts (14) |
| Human | yes | yes | no | 1,008 obese patients (15) |
| Human | yes | no | no | 150 female twins (9) |
| Human | yes | no | no | 449 human donors (8) |
| Human | no | no | yes | Integrated 987 genome-scale data sets (12) |
| Mouse | - | yes | yes | C57BL/6J x A/J mouse cross (16) |
| Mouse | - | yes | yes | C57BL/6J x C3H ApoE -/- mouse cross (17, 18) |
| Mouse | - | yes | yes | C57BL/6J x C3H wildtype mouse cross (19) |
| Mouse | - | yes | yes | C57BL/6J x BTBR Lepob mouse cross (20) |

**Table S3.**
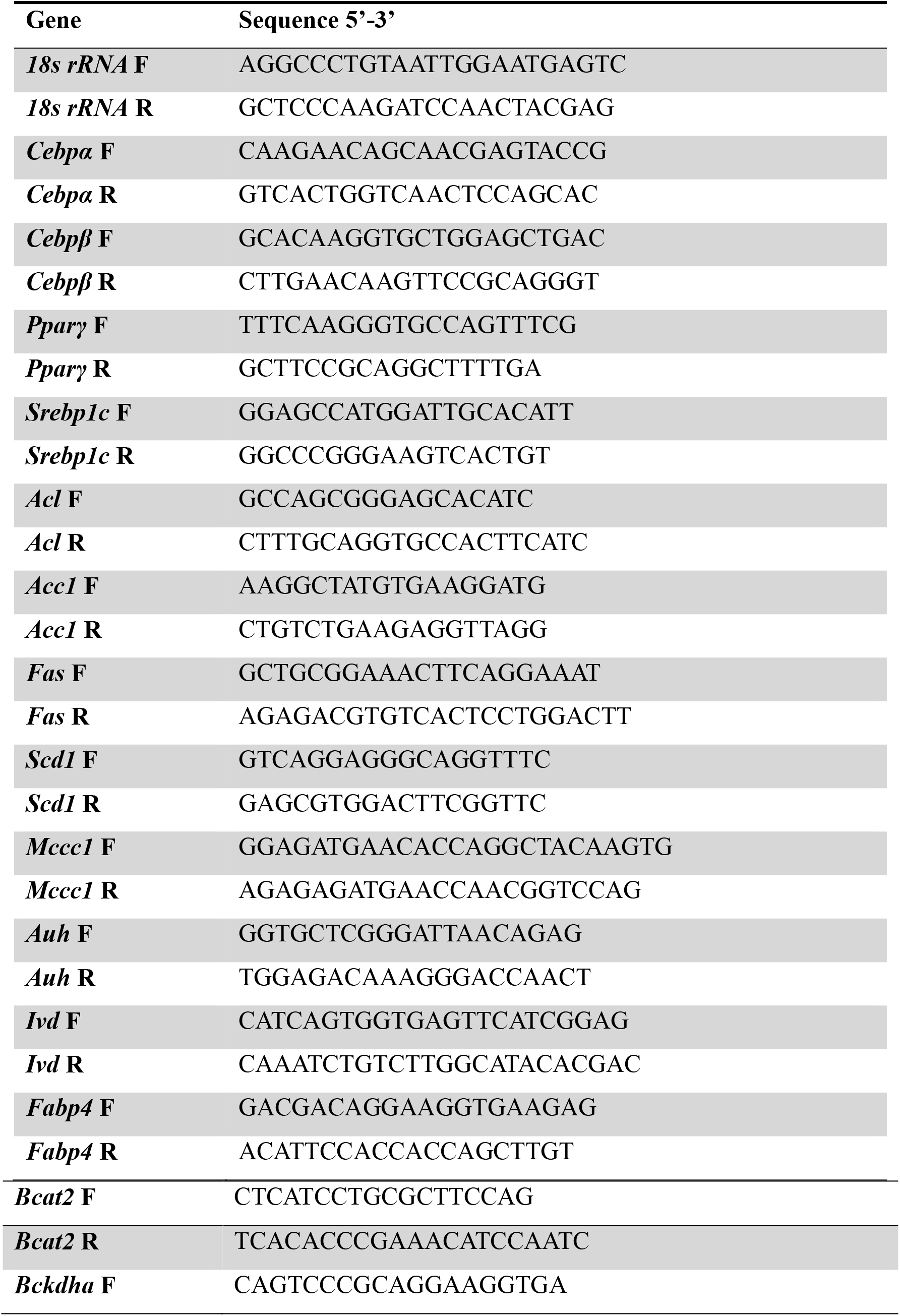
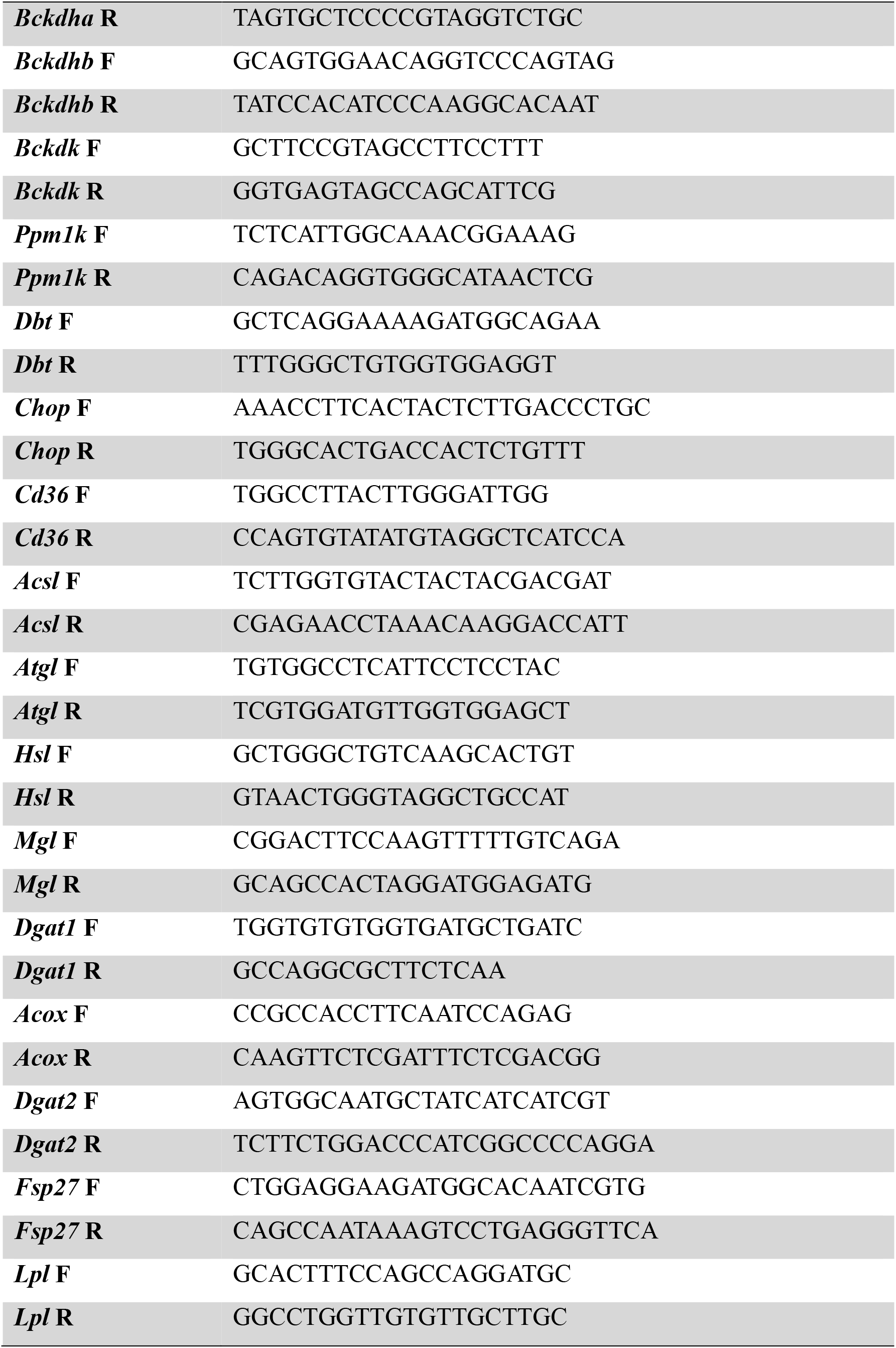
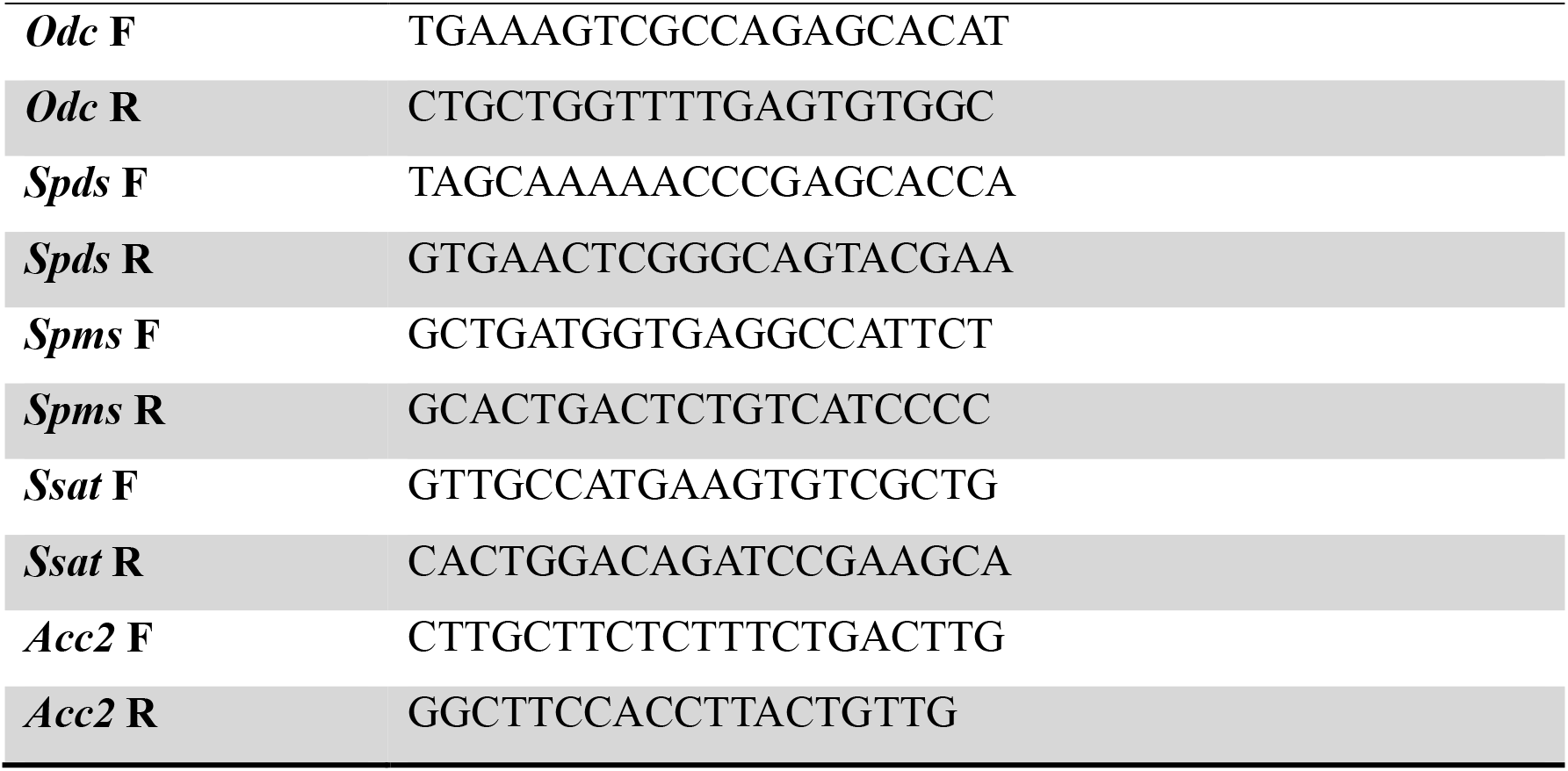
qPCR primers.

